# Memory stabilizes complex ecological systems but delays full restoration

**DOI:** 10.64898/2026.06.05.730340

**Authors:** Yuguang Yang, Serguei Saavedra, Aming Li

## Abstract

Memory effects—defined as the capacity of system states to exert long-lasting influence on subsequent dynamics—are widely recognized as central features of complex living systems. In ecological systems, however, their consequences for stability and recovery dynamics remain poorly understood. To fill this gap, we develop a general theoretical framework that incorporates memory into the dynamics of species-rich ecological systems with complex interaction structures. Our analyses reveal that memory effects expand the stability domain, enabling systems that would otherwise be unstable to persist following perturbations, particularly in cases where instability involves oscillatory behavior. At the same time, memory can accelerate short-term recovery, allowing systems to return more rapidly toward equilibrium in the early stages after perturbation. These apparent benefits, however, come at a cost: memory effects markedly slow long-term recovery, thereby delaying full restoration, as memory retains the influence of past perturbations and hinders a full return to equilibrium. We further support these results by integrating empirical data into the framework. Together, these results reveal fundamental trade-offs mediated by memory—enhanced stability and faster short-term recovery at the expense of delayed full restoration—highlighting the dual role of memory in shaping resilience in complex ecological systems and, more broadly, complex living systems.

## I. INTRODUCTION

The ability of ecological systems to withstand and recover from external perturbations is a key determinant of their persistence. This ability is commonly formalized through stability, which governs whether a system returns to equilibrium following disturbances or instead diverges toward alternative states or extinction [1, 2]. Since the seminal work of May [3], understanding the stability of complex ecological systems has remained a foundational problem, motivating extensive theoretical and empirical efforts to identify how interaction structure, diversity, and nonlinearity shape stability in species-rich systems [4–9].

Most existing theoretical studies implicitly assume that ecological dynamics are memoryless and thus purely state-dependent, such that system evolution is determined solely by the current configuration of species abundances, independent of the trajectory by which that configuration was reached [4, 5]. While this framework has enabled substantial analytical progress, it neglects the possibility that past system states may exert persistent influences on present dynamics.

In many ecological systems, such history dependence is unavoidable. Environmental stressors such as droughts, temperature anomalies, or chemical pollutants can induce physiological, behavioral, or interaction-level changes that persist long after the stressor has been removed. These persistent modifications—ranging from altered metabolic rates to changes in interaction strengths or resource-use strategies—embed information about past conditions into current dynamics [10–13]. As a result, system behavior becomes path-dependent rather than purely state-dependent, a phenomenon broadly referred to as memory.

Empirical evidence across diverse ecological contexts, including planktonic, coral, plant, and soil microbial systems, demonstrates that memory effects can substantially influence recovery and persistence [10, 12, 14]. In parallel, theoretical work has begun to formalize memory using non-Markovian or fractional-order dynamics [15]. However, these studies have largely focused on low-dimensional or species-poor systems. How memory shapes the stability and recovery of species-rich ecological systems with complex interaction networks remains poorly understood.

Here we develop a general theoretical framework that incorporates memory into the dynamics of complex ecological systems while preserving their key structural features. Using this framework, we show that memory systematically stabilizes ecological systems by enlarging the set of conditions under which equilibria remain stable. At the same time, memory accelerates short-term recovery following perturbations. These apparent benefits, however, come at a cost: long-term recovery is substantially slowed, thereby prolonging the time required for full restoration. Together, these results reveal fundamental trade-offs in complex ecological systems: memory stabilizes these systems and enhances short-term recovery at the expense of delayed full restoration. These trade-offs highlight the dual role of memory in shaping resilience, with implications that extend beyond ecology to complex living systems more broadly.

## II. RESULTS

### A. Modeling framework

In classical studies, the dynamical behavior of a memoryless ecological system composed of *S* species is typically described by the following set of ordinary differential equations [1, 2]

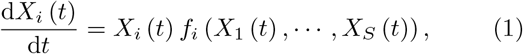

where *X*_*i*_ (*t*) represents the abundance of species *i* at time *t, f*_*i*_ encodes the underlying ecological network (Fig. 1A, B).

**FIG. 1:**
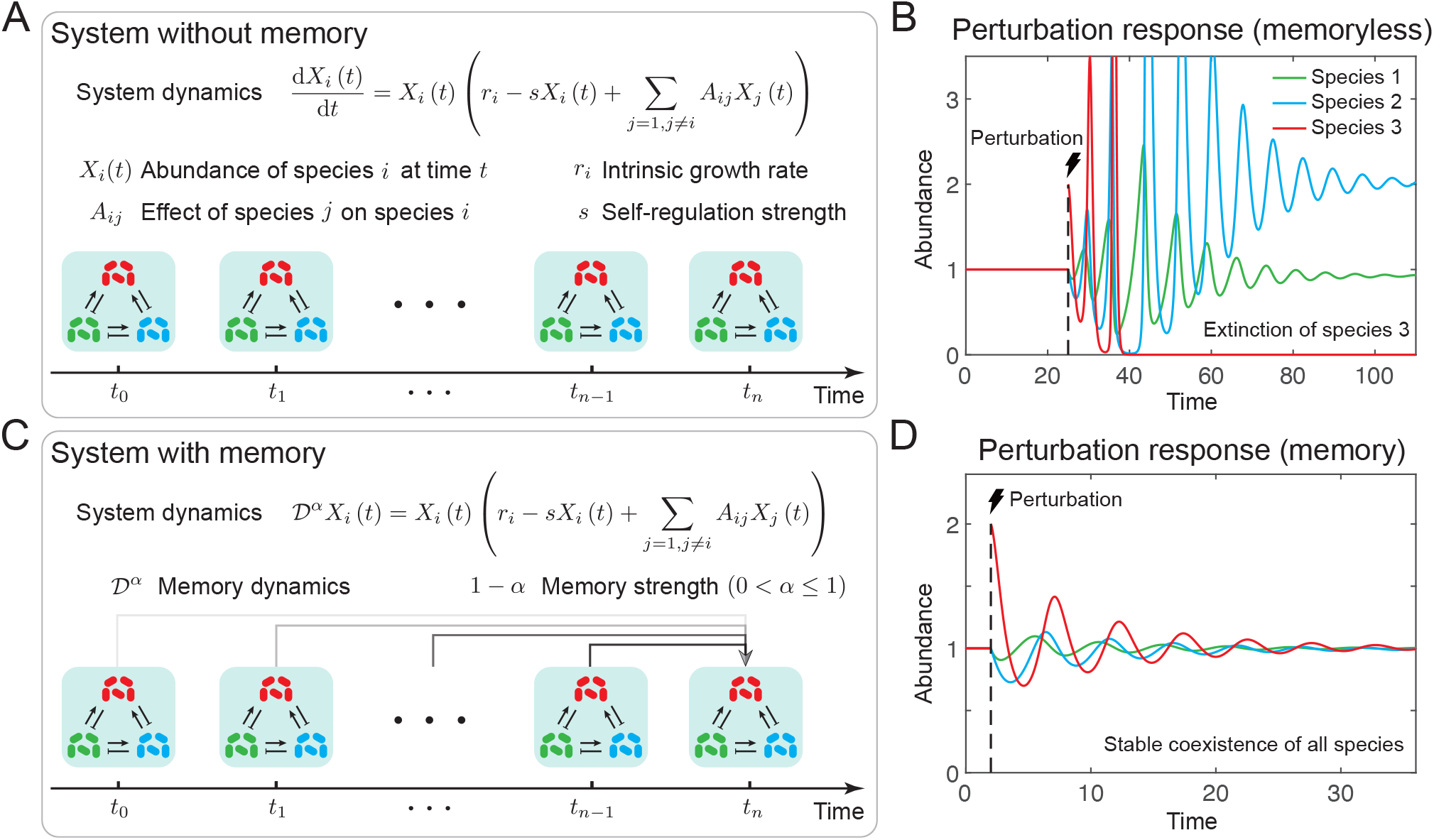
Illustration of system dynamics with and without memory effects. (A) Schematic of a three-species system without memory, whose dynamics follow the generalized Lotka-Volterra model. Colored rods represent different species, with sharp and blunt arrows denoting positive and negative interactions, respectively. In this memoryless case, the systems dynamics are determined solely by the current state. (B) Response of the three-species memoryless system to an external perturbation. After being perturbed, the system fails to return to the original equilibrium, and one species (species 3) goes extinct, indicating that the system is unstable. (C) Schematic of the same three-species system with memory. Here, the system dynamics depend on both the present and all past states. Gray arrows indicate the influence of past states, with lighter arrows representing weaker influence. The system is modeled using fractional-order differential equations instead of traditional ordinary differential equations. Memory strength is quantified as 1 − *α*, where *α* is the fractional derivative order. (D) Response of the three-species system with memory to the same perturbation. The system returns to the original equilibrium, demonstrating that it is stable. This illustrates that memory effects can significantly influence stability. System parameters in panels (B) and (D) are: *r*_1_ = 0.45, *r*_2_ = −1.75, *r*_3_ = −3.95, *s*_*i*_ = 0.05, *A*_12_ = −0.2, *A*_13_ = −0.2, *A*_21_ = 2, *A*_23_ = −0.2, *A*_31_ = 2, *A*_32_ = 2. Memory strength ℳ in panel (D) is 0.15 (i.e., *α* = 0.85).

To incorporate memory effects into system dynamics, we replace the classical first-order derivative operator d /d*t* with the Caputo-type fractional derivative operator 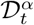, where *α* ∈ (0, 1] denotes the order of differentiation [15–17] (Fig. 1C, D). The Caputo-type fractional derivative is defined via a convolution integral with a power-law memory kernel [16, 17], implying a power-law dependence of current system dynamics on its past states, thereby introducing memory effects into the system (SI Appendix, section 1 and Fig. S1). Accordingly, the governing equations for systems with memory take the form

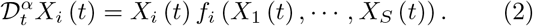

The strength of memory, denoted by ℳ, can be quantified as ℳ = 1 − *α*, such that smaller values of *α* correspond to higher levels of memory [15]. It is worth noting that when *α* = 1, the fractional model reduces to the classical memoryless formulation given by Eq. (1).

If an equilibrium 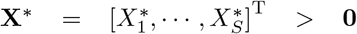 (i.e., all species have positive abundances) satisfies 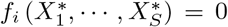 for all *i*, then it is referred to as a feasible equilibrium [18–20]. Ecologists are particularly interested in analyzing system dynamics around such equilibria, as the emergence of an unfeasible equilibrium implies the extinction of one or more species. The dynamical behavior around a feasible equilibrium can be approximated by linearization, leading to

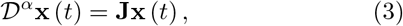

where **x** (*t*) = **X** (*t*) **X**^*^ is the deviation from the equilibrium, and **J** is the Jacobian matrix evaluated at the equilibrium (the so-called “community matrix”) whose element *J*_*ij*_ depicts the effect of species *j* on species *i* near the equilibrium. The stability of the feasible equilibrium can be determined by analyzing the eigenvalues *λ* of the community matrix **J**. For memoryless systems [1, 2], the classical stability criterion requires that all eigenvalues have negative real parts (i.e., Re (*λ*) *<* 0, which is equivalent to | arg (*λ*) | *> π* /2, where arg (*λ*) denotes the angle between the positive real axis and the vector representing *λ* in the complex plane). In contrast, for systems with memory [16, 17], the stability criterion becomes | arg (*λ*) | *> απ* /2 (Methods and SI Appendix, section 1).

Following the tradition initiated by May [3], we model ecological systems by directly constructing the community matrix **J**. In this study, we focus primarily on exploitative systems with trophic structure. Such systems are generated using the cascade model [5, 21, 22], in which species are assumed to form a strict hierarchy (i.e., trophic levels), and interactions occur between any pair of species with a fixed probability, with positive effects from lower-to higher-ranked species and negative effects from higher-to lower-ranked species (Methods).

### B. Memory effects expand the stability domain

While analytical stability criteria are well established, expressing them as geometric regions in the complex plane offers a more intuitive perspective on how memory effects influence stability. For memoryless systems, the stability region corresponds to the entire left half of the complex plane, and the instability region occupies the entire right half plane (Fig. 2A). In contrast, for systems with memory, the stability conditions are relaxed: in addition to the memory-independent stability region in the left half plane, two additional memory-induced stability regions emerge in the right half plane (Fig. 2A). As a result, the instability region is reduced to a sector-shaped region in the right half of the complex plane (Fig. 2A). This geometric pattern indicates that memory effects can stabilize many systems that would be unstable under memoryless dynamics, thereby substantially promoting stability (Fig. 2B).

**FIG. 2:**
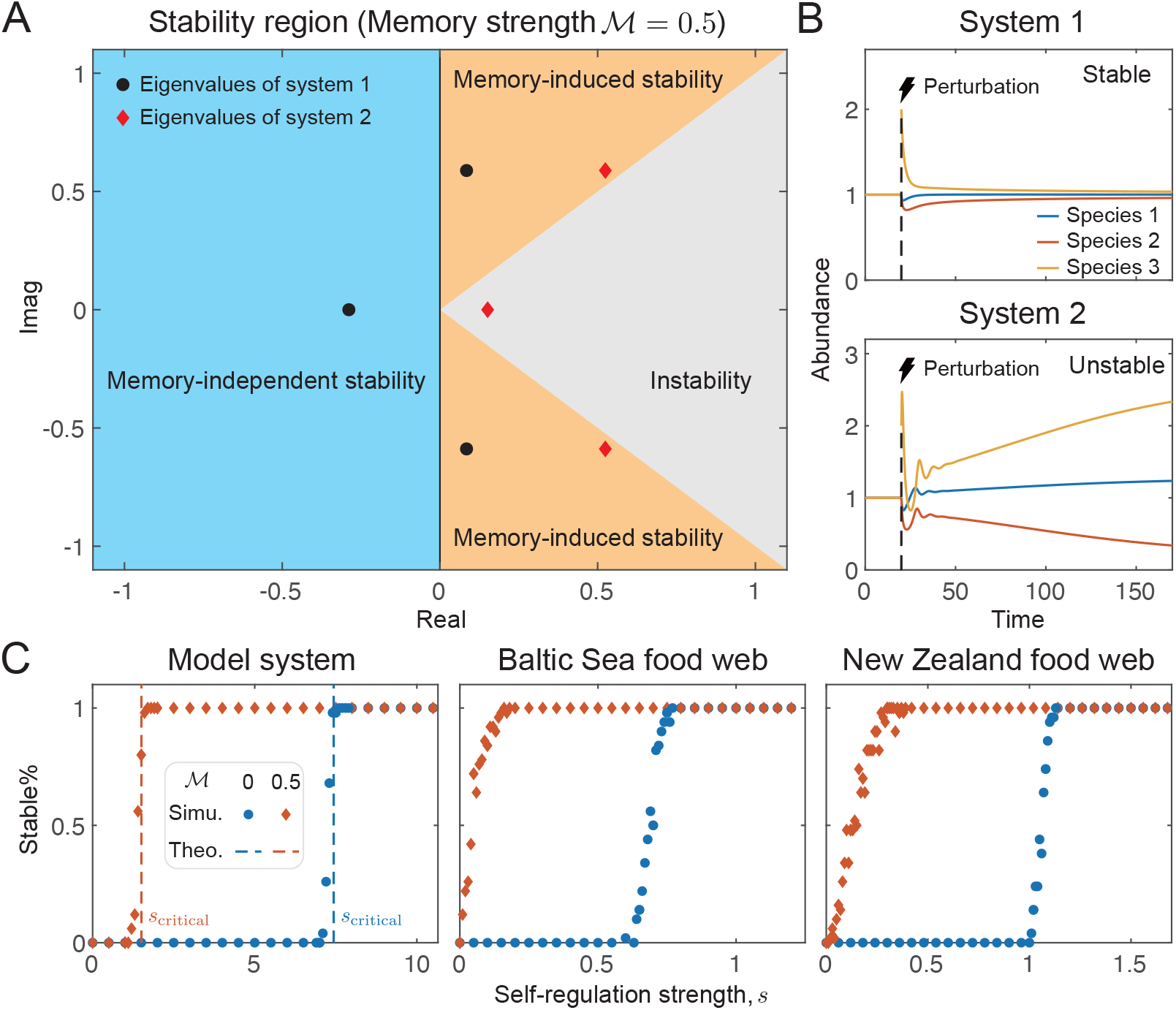
Memory effects stabilize ecological systems. (A) Stability region in the complex plane for systems with memory. Colored regions indicate different stability regimes for the eigenvalues of the community matrix: memory-independent stability (blue), memory-induced stability (orange), and instability (gray). For memoryless systems, stability requires all eigenvalues to lie within the blue region. With memory, stability is ensured if eigenvalues fall within either the blue or orange regions. Shown are eigenvalues of two representative three-species exploitative systems with trophic structure: black dots correspond to system 1, and red diamonds to system 2. In the presence of memory, all eigenvalues of system 1 lie within the stable regions, predicting stability, whereas one eigenvalue of system 2 lies outside these regions, predicting instability. (B) Responses of the two representative systems to an external perturbation. As predicted, system 1 (top) returns to the original equilibrium, indicating stability, while system 2 (bottom) fails to recover, indicating instability. (C) Stability likelihood across varying self-regulation strengths *s* for model and empirical systems. Blue and red dots show simulation results for systems without and with memory, respectively, and dashed lines denote theoretical critical thresholds. Each dot summarizes simulations from 20 independently sampled systems. Memory allows systems to remain stable over a broader range of *s* compared to memoryless systems. The memory strength considered in this figure is ℳ = 0.5. In panel (B), predation and benefit strengths are set to 0.12 and 1.2, respectively. In (C), the model food web uses *S* = 500 and *C* = 0.1; other parameters for model and empirical webs are *µ*_*X*_ = −0.1, *µ*_*Y*_ = 0.9, *σ* = 0.05, and *ρ* = −0.7.

With the stabilizing role of memory effects established, a natural question arises: which specific types of ecological systems are most likely to benefit from it? This question can be addressed by examining how the corresponding memoryless system loses stability. If instability arises due to a real eigenvalue crossing the imaginary axis, then the system cannot be stabilized by memory effects, as the memory-induced stability regions do not include the positive real axis. In contrast, if instability is caused by a pair of complex conjugate eigenvalues crossing the imaginary axis (a signature of Hopf bifurcation [23]), then the system can benefit from memory effects, as the region of the complex plane that was previously unstable for such eigenvalues now becomes part of the stability region.

Indeed, a considerable number of ecological systems loses stability through Hopf bifurcation. Typical examples include predator-prey systems and competitive systems [24–26]. Focusing on the species-rich systems considered in our work, theoretical analysis reveals that the instability of systems under strong positive effects is triggered by a pair of complex conjugate eigenvalues crossing the imaginary axis [5], and can thus be significantly stabilized by memory effects (SI Appendix, section 2 and Figs. S2-S3). As is shown in Fig. 2C, if self-regulation strength is chosen as the control parameter, the critical self-regulation strength for systems with memory to maintain stability is much lower than that for memoryless systems. The same pattern is observed when empirical food web structures are applied (Fig. 2C). Here, a strong positive effect refers to cases where the positive influence from lower-to higher-ranked species outweighs the negative feedback in the reverse direction. Mathematically, for a pair of species *i* and *j*, with interactions described by the symmetric pair (*J*_*ij*_, *J*_*ji*_) of the community matrix **J**, where *J*_*ij*_ *>* 0 and *J*_*ji*_ *<* 0, a strong positive effect occurs when the absolute value of the positive effect exceeds that of the negative one, i.e., | *J*_*ij*_ | *>* | *J*_*ji*_ |. In practice, such interaction patterns are often found in systems with inverted biomass pyramids, typically in planktonic or other aquatic systems [5, 27, 28].

### C. Bounded stabilizing influences of elevating memory strength

Having identified which types of systems benefit from memory, we next ask whether increasing memory strength can continuously enhance the stabilizing influence. To explore this, consider a case in which instability arises due to a decrease in self-regulation strength. In this context, the question becomes: can increasing memory strength continuously lower the critical self-regulation threshold required to maintain stability?

In addressing this question, we first examine how increasing memory strength influences the size of the stability region. We define the stability region size, denoted by *θ*, as the ratio of the stability region to the entire complex plane (a dimensionless geometric measure). Specifically, *θ* is calculated by dividing the sector angle of the stability region by the total angle of the complex plane, which is 2*π*. Since the instability region spans an angle of *απ*, the stability region size can be quantified as *θ* = (1 + ℳ) /2 (with ℳ = 1 − *α*, as introduced above; derivation of this expression is provided in SI Appendix, section 1).

Clearly, *θ* increases monotonically with memory strength (Fig. 3A), suggesting that increasing memory strength may lead to an unbounded stabilizing influence.

**FIG. 3:**
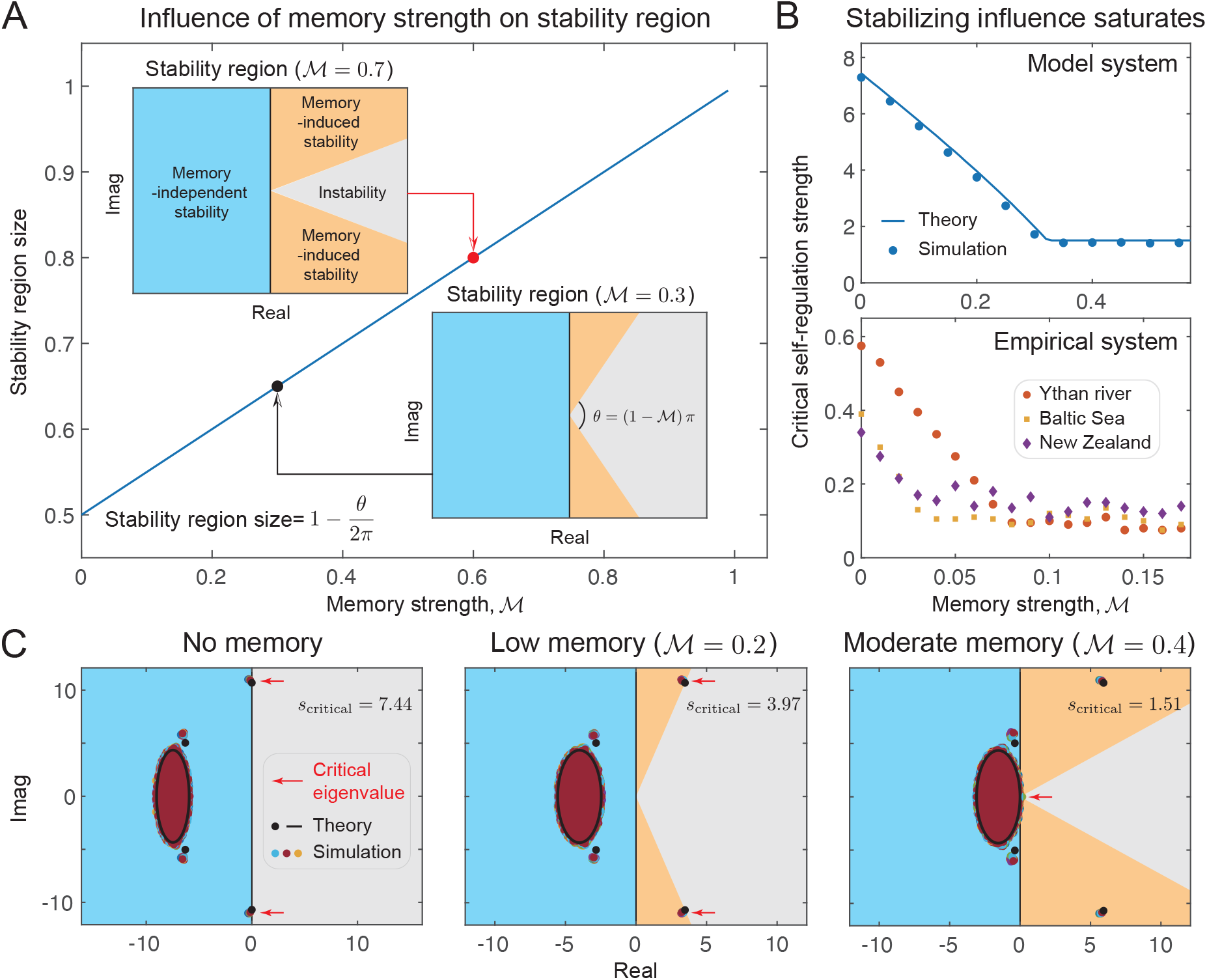
Stabilizing influence of memory effects saturates. (A) Relationship between memory strength ℳ and the size of the stability region. Insets show stability regions in the complex plane for two representative memory strengths. The stability region size is defined as the fraction of the complex plane satisfying the stability condition and is calculated as 1 − *θ* /(2*π*), where *θ* = (1 − ℳ)*π*. As memory strength increases, the stability region expands monotonically. (B) Relationship between memory strength ℳ and the critical self-regulation strength. Top and bottom panels show results for model and empirical systems, respectively. Dots, squares, and diamonds represent results from numerical simulations (each symbol indicates the average of 50 independently sampled systems), while the blue line denotes the theoretical prediction. For both model and empirical systems, increasing memory strength initially lowers the critical self-regulation threshold, but the stabilizing influence plateaus as memory strength continues to grow. (C) Eigenvalue distributions of representative model systems at the critical state under different memory strengths. Colored dots show simulation results; black dots and lines represent theoretical predictions. Red arrows indicate critical eigenvalues that lie at the stability boundary and thus determine system stability. In systems without memory (left panel) or with low memory (middle panel), the critical eigenvalues form a complex conjugate pair. Since the stability region for complex eigenvalues expands with memory, increasing memory in these cases exerts a stabilizing influence. However, in systems with moderate memory (right panel, and also at high memory), the critical eigenvalue is real, and the stability region does not expand for such eigenvalues—thus, further increases in memory yield no additional stabilizing influence. In panels (B) and (C), model systems use *S* = 500 and *C* = 0.1; other parameters for model and empirical systems are *µ*_*X*_ = −0.1, *µ*_*Y*_ = 0.9, *σ* = 0.05, and *ρ* = −0.7.

Surprisingly, however, this influence is in fact bounded (Fig. 3B). As memory strength increases, the critical self-regulation threshold initially decreases, but eventually plateaus once memory exceeds a certain level, indicating that further increases do not enhance stability. Mathematically, this can be explained by the eigenvalue distribution at the critical self-regulation strength (Fig. 3C). When the system has no or low level of memory, a pair of complex conjugate eigenvalues is the first to leave the stability region, so enlarging the stability region for complex eigen-values significantly enhances stability. At moderate (and also high) level of memory, a real eigenvalue becomes the limiting factor; since the stability region does not extend along the positive real axis, further increases in memory strength no longer provide additional stabilizing effects.

### D. Memory effects accelerate short-term recovery

In addition to evaluating how memory effects influence stability—that is, whether a system can eventually recover from external perturbations—it is equally important to investigate the detailed dynamics of the recovery process itself. Ecologists are particularly interested in understanding the speed of recovery and often distinguish between two key dimensions: short-term and long-term recovery [29–33]. This raises an important question: how do memory effects shape these two aspects of recovery dynamics? To address this, we need to first specify how recovery performance is quantified. In this study, recovery performance is evaluated by measuring the distance *D* between the current system state and the equilibrium state at a fixed time *T*_perturbed_ following the perturbation. Specifically, if the perturbation occurs at time *t*_0_, the distance is assessed at *t*_0_ + *T*_perturbed_.

We first focus on short-term recovery. Here, we consider systems in which all perturbations decay initially (i.e., non-reactive systems) for convenience, as such systems can in principle exhibit monotonic responses to perturbations [31–33] (the distance to equilibrium decreases continuously following a perturbation). For short-term recovery, a relatively small fixed time *T*_perturbed_ is chosen, and the corresponding distance *D* is denoted as *D*_short_. Our results show that *D*_short_ for systems with memory is smaller than that for their memoryless counterparts, suggesting that short-term recovery is accelerated by memory effects (Fig. 4A). Theoretical analysis of initial recovery rate also supports this finding (Methods, SI Appendix, section 3 and Fig. S4). Similar to our analysis of stabilizing influence, a natural question arises: does increasing memory strength continuously accelerate short-term recovery? Further numerical simulations of both model and empirical systems confirm that the answer is yes. Specifically, increasing memory strength leads to a continuous reduction in *D*_short_, indicating that the degree of short-term recovery enhancement grows with memory strength (Fig. 4B).

**FIG. 4:**
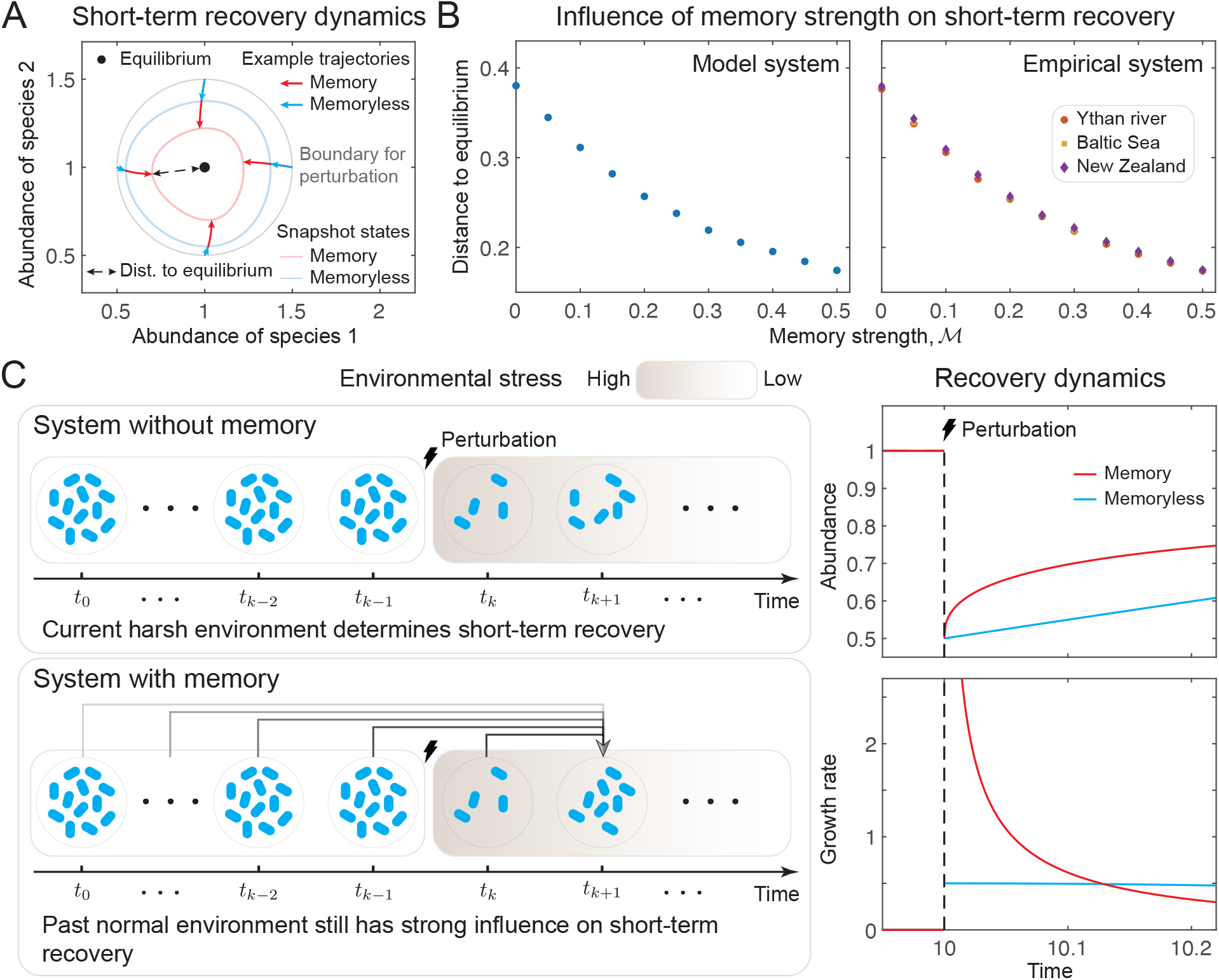
Memory effects accelerate short-term recovery. (A) Snapshots of short-term recovery dynamics for systems without and with memory. One hundred equal-amplitude perturbations, uniformly distributed across all directions, are applied at *t*_0_, and system states are recorded at *t*_0_ + 0.1. Light blue and red curves represent the collections of snapshot states for systems without and with memory, respectively, while dark blue and red arrows show four representative trajectories. The black dot marks the equilibrium, and the dashed arrow indicates the distance from the snapshot state to the equilibrium. In all cases, trajectories of systems with memory end closer to equilibrium, indicating faster short-term recovery. (B) Influence of memory strength on short-term recovery in model (left) and empirical (right) systems. Recovery performance is quantified by the Euclidean distance between the post-perturbed state and the equilibrium at a fixed post-perturbation time *T*_perturbed_ = 0.1 (i.e., dashed arrow in panel (A)). Each data point represents the mean of 50 independently sampled systems. Larger memory strength consistently yields improved short-term recovery (i.e., smaller distances). (C) Conceptual toy model illustrating the intuition by which memory enhances short-term recovery. A single population has a high (low) growth rate under low (high) environmental stress. A perturbation reduces both population size and environmental quality is applied. In the short term, the memoryless population (top left) responds solely to the current harsh conditions, exhibiting a low growth rate and thus slow recovery (blue curves, right). In contrast, the population with memory (bottom left) “remembers” influence from prior favorable conditions, maintaining a higher growth rate and recovering more rapidly (red curves, right). In panel (A), simulations are based on generalized Lotka-Volterra model with parameters *r*_1_ = 2.5, *r*_2_ = 1.5, *A*_12_ = −0.5, *A*_21_ = 0.5, *A*_11_ = *A*_22_ = −2, and ℳ = 0.6. In panel (B), model systems use *S* = 100, *C* = 0.1; for both model and empirical systems, other parameters are *s* = 10, *µ*_*X*_ = −0.1, *µ*_*Y*_ = 0.9, *σ* = 0.05, and *ρ* = −0.7. In panel (C), simulations are based on the fractional-order logistic model with parameters *s* = 2 and ℳ = 0.6.

This enhancing influence can be intuitively understood using a conceptual toy model of a single population (Fig. 4C). We assume that the population exhibits a relatively high growth rate under favorable environmental conditions (i.e., low environmental stress) and a relatively low growth rate under unfavorable conditions (i.e., high environmental stress). Before the perturbation is applied, the population is assumed to have already reached its equilibrium state. Following the perturbation—which simultaneously reduces population size and increases environmental stress—the responses of populations with and without memory diverge. In the short term after the perturbation, the memoryless population responds immediately to the elevated environmental stress, resulting in a low growth rate and thus slow short-term recovery. In contrast, the population with memory retains a “memory” of the previously favorable environmental conditions and temporarily maintains a higher growth rate, leading to faster short-term recovery.

Finally, although our main analysis of short-term recovery focuses on non-reactive systems, accelerated short-term recovery due to memory effects can also be observed in reactive systems (SI Appendix, Fig. S5).

### E. Memory effects inhibit long-term recovery

We now turn to long-term recovery. Indeed, long-term recovery, commonly studied through the concept of “resilience”, has been a central and long-standing focus in ecology [1, 2, 29, 30, 34]. Typically, long-term recovery is quantified by the asymptotic recovery rate, which captures how rapidly a system returns to equilibrium following perturbations once transient dynamics have decayed and thus reflects the time required for full restoration [29, 30]. Long-term recovery is widely regarded as closely linked to stability: systems that require less time for full restoration are often viewed as more stable, whereas systems that require more time tend to be less so, and instability implies that full restoration cannot be achieved. Moreover, stabilizing factors are often thought to simultaneously promote long-term recovery. For example, classic studies have demonstrated that increasing self-regulation strength can both stabilize an otherwise unstable system and enhance long-term recovery in an already stable one [4, 8].

Building on these insights, we next investigate how memory effects influence long-term recovery. To assess this, we select a relatively large fixed time *T*_perturbed_ and denote the corresponding distance to equilibrium as *D*_long_. Given that previous analyses have demonstrated the stabilizing role of memory effects—and that such effects enhance short-term recovery—one might reasonably expect memory to also improve long-term recovery. Surprisingly, this expectation does not hold: memory effects significantly inhibit long-term recovery (Fig. 5A, Methods and SI Appendix, section 3). Our results show that *D*_long_ for systems with memory is consistently greater than that of their memoryless counterparts. Furthermore, theoretical analysis and numerical simulations reveal that long-term recovery performance deteriorates further as memory strength increases (Methods, SI Appendix, section 3 and Fig. S4). This trend is consistently observed in both model and empirical systems (Fig. 5B).

**FIG. 5:**
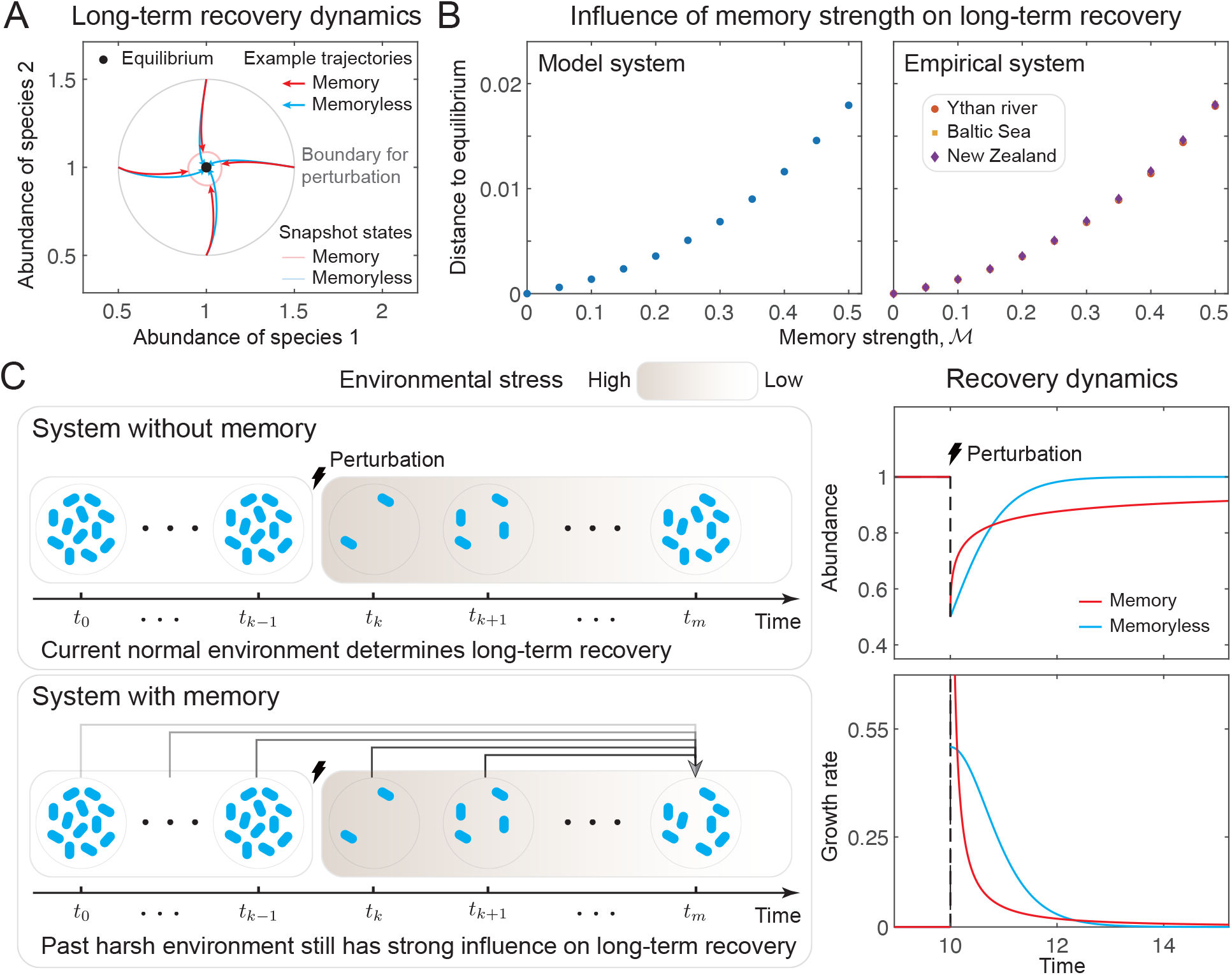
Memory effects slow down long-term recovery. (A) Snapshots of short-term recovery dynamics for systems without and with memory. One hundred equal-amplitude perturbations, uniformly distributed across all directions, are applied at *t*_0_, and system states are recorded at *t*_0_ + 3. Light blue and red curves represent the collections of snapshot states for systems without and with memory, respectively, while dark blue and red arrows show four representative trajectories. The black dot marks the equilibrium. In all cases, trajectories of systems with memory end farther from the equilibrium, indicating slower long-term recovery. (B) Influence of memory strength ℳon long-term recovery in model (left) and empirical (right) systems. Long-term recovery performance is quantified by the Euclidean distance between the perturbed state and the equilibrium at a fixed post-perturbation time *T*_perturbed_ = 10. Each data point represents the mean of 50 independently sampled systems. As memory strength increases, long-term recovery performance consistently deteriorates (i.e., distance increases). (C) Conceptual toy model illustrating the intuition by which memory inhibits long-term recovery. A single population has a high (low) growth rate under low (high) environmental stress. A perturbation reduces both population size and increases environmental stress is applied. In the long term, the memoryless population (top left) responds to the current benign conditions and recovers efficiently (blue curves, right). In contrast, the population with memory (bottom left) continues to retain influence from the earlier harsh environment, resulting in a suppressed recovery rate and slower recovery (red curves, right). The system parameters are the same as those used in Fig. 4.

Although unexpected, this inhibiting influence can once again be intuitively understood through the conceptual toy model introduced earlier (Fig. 5C). As before, we assume that a perturbation simultaneously reduces population size and increases environmental stress. In the long term following the perturbation, environmental conditions are assumed to have returned to—or at least approached—their original, favorable state. At this stage, the memoryless population responds directly to the improved environment, exhibiting a relatively high growth rate that facilitates faster long-term recovery. In contrast, the population with memory preserves a “memory” of the previously elevated environmental stress, which continues to suppress its growth rate and thereby leads to slower long-term recovery.

## III. DISCUSSION

Since the pioneering work by May in 1972, substantial progress has been made in understanding the drivers of stability and instability in complex ecological systems [3–9, 35–42]. However, most of these advances assume that system dynamics are purely state-dependent, neglecting memory effects—i.e., it is path-dependent, with present behavior shaped not only by the state at a specific instant but also by the trajectory leading to it—which are widely observed in the real world. As a result, our understanding of stability and recovery dynamics has been constrained by this oversight. In this study, we have sought to disentangle the role of memory effects in shaping ecological dynamics. Through analyses of both model and empirical systems, we demonstrate that memory effects act as a double-edged sword in regulating stability and recovery dynamics. While memory can significantly enhance both stability and short-term recovery, it concurrently impairs long-term recovery. We also consider different species abundance distributions and find that they do not alter the main conclusions (SI Appendix, section 4 and Figs. S6-S8). These findings suggest that ecological systems may face two fundamental and consequential trade-offs: one between stability and the time required for full restoration, and another between short-term and long-term recovery (Fig. 6).

**FIG. 6:**
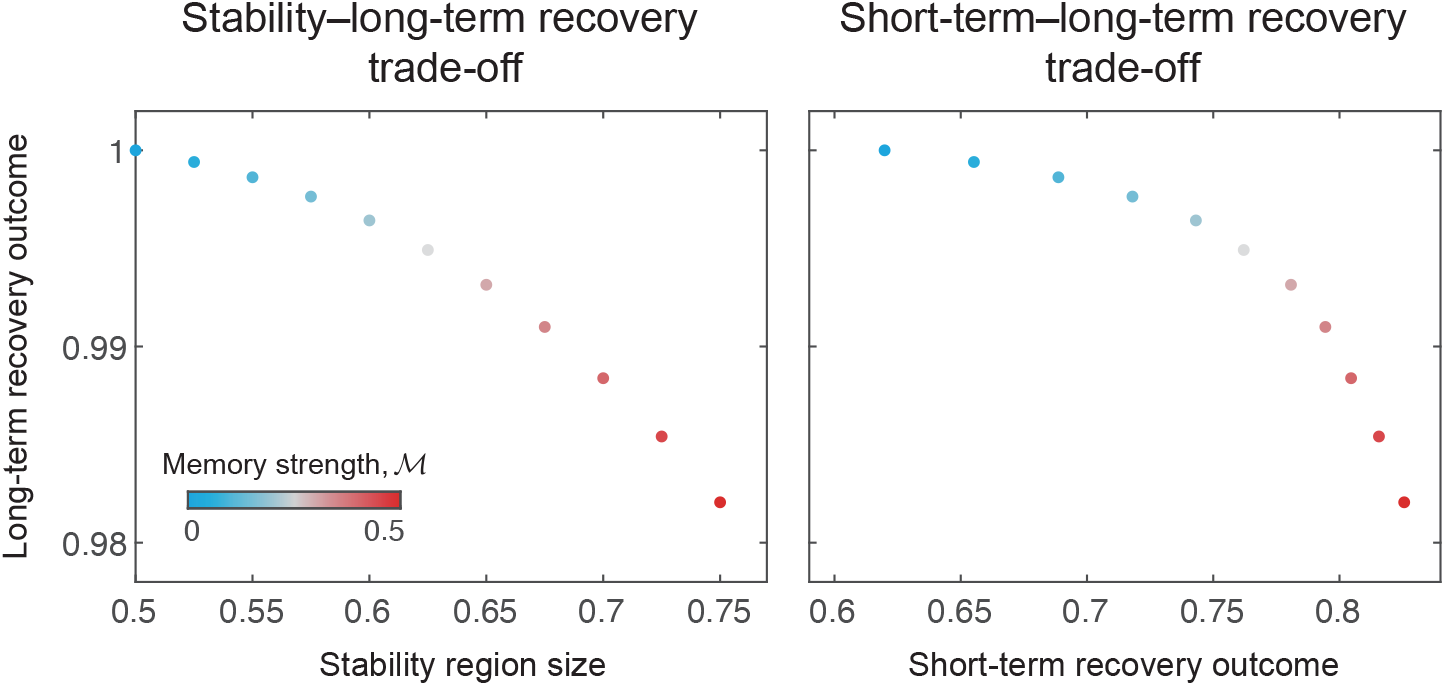
Memory-induced trade-offs in complex ecological systems. blueLong-term recovery outcome is shown against stability region size (left panel) and short-term recovery outcome (right panel). Each data point represents the mean over 50 independently sampled systems, with color indicating memory strength. Increasing memory strength expands the stability domain (left) and enhances short-term recovery (right), but substantially impedes long-term recovery (both panels). These patterns illustrate two fundamental trade-offs induced by memory: one between stability and the time required for full restoration (left), and the other between short-term and long-term recovery (right). Recovery outcome is defined as the fraction of the perturbation that is recovered, measured as the recovered distance relative to the total displacement. Short-term recovery outcome is measured at post-perturbation time *T*_perturbed_ = 0.1, while long-term recovery outcome is measured at *T*_perturbed_ = 10. System parameters are the same as in Fig. 4.

The first trade-off implies that, in order to maintain stability—or to preserve a high stability margin in the face of environmental fluctuations—ecological systems may need to sacrifice the time required for full restoration following perturbations. This leads to an interesting implication: systems that require more time to recover may in fact, be those better equipped to retain stability under variable conditions. Moreover, increasing recovery time (i.e., critical slowing down) is widely regarded as an early-warning signal of instability and impending critical transitions [43]. However, this trade-off suggests that this belief should be revisited. Increasing memory strength improves stability while simultaneously diminishing long-term recovery, whereas decreasing memory strength impairs stability but enhances long-term recovery. This inversion highlights the need to reassess how critical slowing down indicators are interpreted in systems with memory.

The second trade-off highlights a temporal tension between short-term and long-term recovery. Memory effects enhance short-term recovery by preserving beneficial information from prior states, allowing systems to rebound more quickly in the early phases following perturbations. However, this same mechanism impedes long-term recovery, as the retained memory of past stress continues to influence system dynamics even after environmental conditions have normalized. As a result, systems with stronger memory exhibit faster initial responses but slower eventual convergence to equilibrium. This trade-off implies that ecological systems cannot simultaneously optimize both rapid early recovery and efficient long-term adjustment. In practical terms, systems tuned for short-term robustness— such as those frequently exposed to recurring, transient perturbations—may inherently compromise their capacity for full recovery. Conversely, systems that prioritize long-term recovery may experience sluggish short-term responses. This duality offers an important perspective for interpreting ecological recovery trajectories and designing restoration strategies, especially in environments where the frequency and duration of perturbations vary widely.

Taken together, our study presents a formal theoretical framework that can be integrated with empirical data to advance understanding of stability and recovery dynamics in ecological systems exhibiting memory effects, a characteristic feature of complex living systems. Using this framework, we demonstrate how memory regulates stability and recovery dynamics and uncover two fundamental trade-offs faced by ecological systems. These findings provide informative guidance for predicting ecological responses and for informing the design and implementation of control and intervention strategies in complex ecological systems and complex living systems. Moreover, although our analysis is framed in ecological terms, the results are broadly applicable to other complex systems, as they are rooted in general principles of dynamical systems theory.

## IV. METHODS

### A. Stability analysis of systems with memory

Dynamical systems theory [23] suggests that a feasible equilibrium is locally asymptotically stable (hereafter, stable) if lim_*t*→+∞_ **x**(*t*) = **0** for any sufficiently small perturbation (**x** (*t*) represents the deviation from the equilibrium). Since around the feasible equilibrium, the dynamical behavior can be approximated by the linearized system (i.e., Eq. (3)). Stability can thus be determined by analyzing this linearized system. For memoryless systems, the solution to the linearized system can be expressed as

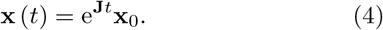

Here **x**_0_ is the perturbation applied at time *t*_0_. To ensure lim_*t*→+∞_ **x** (*t*) = **0**, all eigenvalues of the community matrix **J** should have negative real parts. That is, the stability criterion for memoryless systems reads as Re(*λ*) *<* 0 for all eigenvalues *λ*—or equivalently, | arg(*λ*) | *> π/*2. For systems with memory, the solution to the linearized system can be expressed as

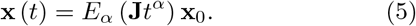

Here *E*_*α*_ (·) is the Mittag-Leffler function [44], and is defined as

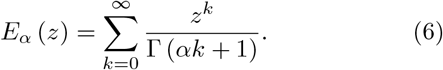

According to the asymptotic properties of this Mittag-Leffler function [44], to ensure lim_*t*→+∞_ **x** (*t*) = **0**, all eigenvalues *λ* of the community matrix **J** should satisfy |arg(*λ*)| *> απ/*2. That is, the stability criterion for systems with memory reads as |arg(*λ*)| *> απ/*2.

### B. Numerical implementation of fractional-order dynamics

Fractional-order differential equations in this study were numerically solved using the `fde12` solver, a publicly available MATLAB implementation developed by Garrappa [45]. The `fde12` algorithm is based on the predictor–corrector scheme of the Adams–Bashforth–Moulton (ABM) type for Caputo fractional derivatives. In this method, the solution at each time step is first predicted using an explicit Adams–Bashforth formula and subsequently corrected using an implicit Adams–Moulton step, ensuring both stability and accuracy for fractional orders *α* ∈ (0, 1). The local truncation error of the method scales as 𝒪 (*h*^1+*α*^), where *h* denotes the integration step size. All numerical integrations were performed with sufficiently small step sizes to ensure numerical convergence and stability.

### C. Construction of the community matrix

In our main analyses, ecological systems are modeled by directly constructing their community matrices. For model systems, we follow the widely used cascade model, and the community matrix is generated through the following procedure [4, 5, 21]. First, for each off-diagonal entry pair (*J*_*ij*_, *J*_*ji*_)_*i* ≠*j*_, we draw a random number *p* from a uniform distribution *U* [0, 1]. If *p > C*, the pair is set as (*J*_*ij*_, *J*_*ji*_) = (0, 0), where *C* denotes the connectance of the community and controls the density of species interactions. If *p* ≤ *C*, the pair (*J*_*ij*_, *J*_*ji*_) is sampled from a bivariate distribution *Z* = (*X, Y*) with mean vector ***µ*** and covariance matrix **Σ** given by

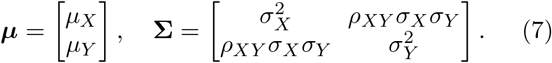

The marginal distribution *X* characterizes the effects of higher-ranked species on lower-ranked species, and hence *µ*_*X*_ *<* 0, while *Y* describes the effects of lower-ranked species on higher-ranked species, with *µ*_*Y*_ *>* 0. For simplicity, we assume *σ*_*X*_ = *σ*_*Y*_ = *σ*. Finally, all diagonal elements *J*_*ii*_ are set to *−s*, indicating that all species self-regulate with the same strength *s*. For systems with empirical food-web structures, the original datasets provide the binary adjacency matrix that specifies whether an interaction exists between any pair of species. For each interacting pair, the corresponding interaction strengths (*J*_*ij*_, *J*_*ji*_) are sampled from the same bivariate distribution *Z* = (*X, Y*), whose mean vector ***µ*** and covariance matrix **Σ** are specified in Eq. (7).

### D. Eigenvalue distribution of the cascade community matrix

Recent advances in random matrix theory enable the derivation of the eigenvalue distribution of the cascade community matrix [5]. For convenience, the following parameters are defined

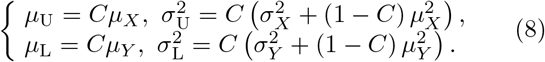

To obtain the eigenvalue distribution of the community matrix **J**, we consider the eigenvalues of the shifted matrix 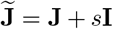, since the eigenvalues of **J** can be recovered by translating those of 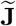 leftward by *s* units. The matrix 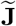 is decomposed as 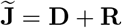, where **D** is a deterministic matrix whose upper triangular entries are all equal to *µ*_U_ and lower triangular entries are all equal to *µ*_L_. The residual matrix is defined as 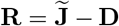. The eigenvalues of 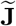 can be approximated by combining the eigenvalue distributions of **D** and **R**. The eigenvalues of **D** lie on a circle and can be expressed as

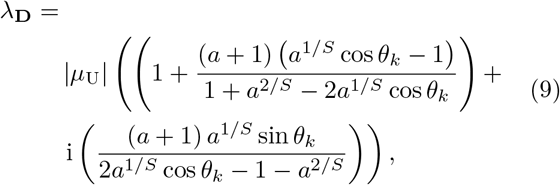

where

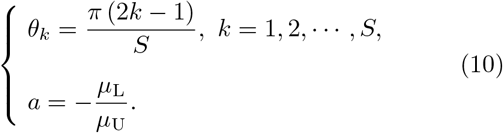

The eigenvalues of **R** are confined within an ellipse (SI Appendix, section 2), defined by

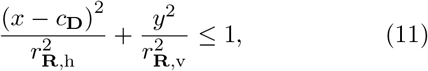

where

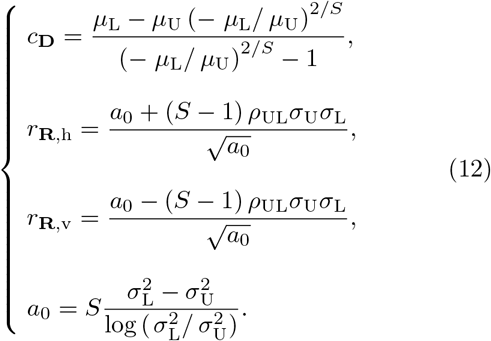

By combining the eigenvalue distributions of **D** and **R**, we obtain an approximation of the eigenvalue distribution of 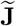. The eigenvalue distribution of **J** is then recovered by incorporating the effect of the diagonal self-regulation terms.

### E. Initial recovery rates for systems with and without memory

In analyzing recovery rates, we assume that both systems with memory and their memoryless counterparts are stable. For such systems resting at a feasible equilibrium **X**^*^, the (average) recovery rate *r*(*t*) is defined as [29, 30]

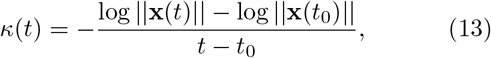

where **x**(*t*) = **X**(*t*) − **X**^*^ denotes the deviation from equilibrium, and ||·|| is the 2-norm.

As noted above, for the analysis of short-term recovery (and initial recovery rate), we focus on non-reactive systems—both memoryless and with memory—since these exhibit a monotonic response to perturbations. The initial recovery rate is defined immediately after the perturbation, i.e., in the limit *t* → 0^+^:

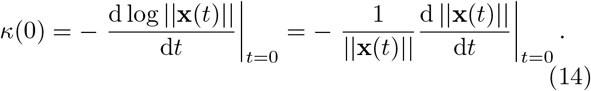

For memoryless systems, this reduces to

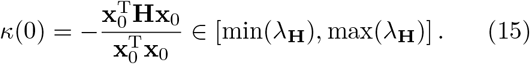

Since we are considering non-reactive systems, max(*λ*_**H**_) *<* 0, implying that the initial recovery rate is a positive finite value [31–33]. For systems with memory, the initial recovery rate is given by

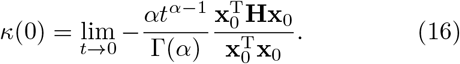

This limit diverges (*κ*(0) = ∞), indicating that systems with memory exhibit substantially higher initial recovery rates than their memoryless counterparts.

In numerical implementations, the initial recovery rate is estimated by evaluating *κ*(*t*) at a small but finite time *t*_*ε*_ *>* 0, rather than taking the singular limit *t* → 0. This ensures that *κ*(0) remains finite and numerically well-defined.

### F. Long-term recovery rates for systems with and without memory

The long-term recovery behavior of stable systems is examined for sufficiently large *t*. For memoryless systems, the long-term (asymptotic) recovery rate can be approximated as [29, 30]

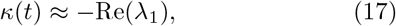

where *λ*_1_ is the rightmost eigenvalue of the community matrix **J**. Since the systems under consideration are stable (i.e., Re(*λ*_1_) *<* 0), this recovery rate remains a positive constant. In contrast, for systems with memory, the long-term recovery rate can be approximated as

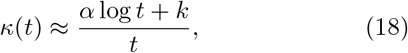

where *k* is a constant. As *t* → ∞, the recovery rate of memoryless systems remains constant, while that of systems with memory gradually declines toward zero. This indicates that, at sufficiently large times, systems with memory exhibit lower long-term recovery rates than their memoryless counterparts.

### G. Empirical food webs

We analyzed three large, published empirical food webs: (i) the food web of the Ythan River estuary on the east coast of Scotland [46], (ii) the food web of a brackish shallow-water inlet in the Baltic Sea [47], and (iii) the food web of an intertidal mudflat ecosystem in New Zealand.[48]. For each empirical food web, we first obtained its topological structure and extracted the adjacency matrix following two steps: (1) self-loops (i.e., cannibalism) were removed; (2) in the rare cases of bidirectional links (i.e., species *a* preys on species *b* and species *b* preys on species *a*), one of the two links was removed at random. Using the resulting adjacency matrix, the corresponding community matrix was then constructed as described above.

## V. ACKNOWLEDGMENTS

We gratefully acknowledge the support from the National Natural Science Foundation of China under Grant Nos. T2525017, 62533002, T2421004, and 62173004, the National Key Research and Development Program of China under Grant No. 2022YFA1008400, and the Jiangsu Provincial Scientific Research Center of Applied Mathematics under Grant No. BK20233002. S.S. acknowledges support from the National Science Foundation under Grant No. DEB-2436069.

## Appendix A: Dynamical framework

### 1. Stability analysis of systems with and without memory

The dynamical behavior of a classical memoryless system (i.e., where current dynamics depend only on the current state) composed of *S* interacting species is described by the following ordinary differential equations (ODEs):

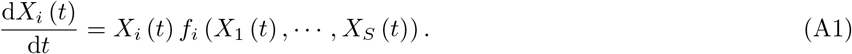

Here, *X*_*i*_ (*t*) denotes the abundance of species *i* at time *t*, and *f*_*i*_ is an unspecified function encoding the ecological interactions.

Memory effects imply that present system dynamics are shaped not only by the current state but also by past configurations. These effects can be modeled by introducing fractional derivatives into the governing equations [15–17]. Accordingly, the behavior of a system with memory can no longer be captured by ODEs (Eq. (A1)) but must instead be described by a set of fractional-order differential equations (FODEs):

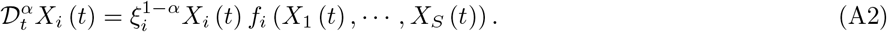

Here, 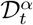 denotes the fractional derivative with order *α*, and *ξ*_*i*_ is the characteristic time scale of species *i*. The parameters *ξ*_*i*_ ensure dimensional consistency between both sides of the equation. For simplicity and analytical convenience, we assume *ξ*_*i*_ = 1 for all species, which reduces Eq. (A2) to

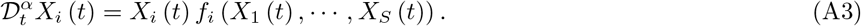

To capture memory effects [15], the order of differentiation *α* must fall within the range (0, 1]. This fractional order quantifies memory strength: when *α* = 1, the equation reduces to the standard ODE, implying no memory; as *α* decreases, the influence of past states increases. Thus, memory strength ℳ can be defined as [15]

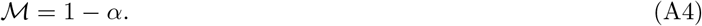

Several definitions of fractional derivatives exist, including Riemann-Liouville, Grünwald-Letnikov, and Caputo types [15–17]. Among them, the Caputo derivative is preferred for real-world systems because it introduces a power-law decay of past influence, a feature commonly observed in nature. In addition, it accommodates classical initial conditions with well-defined physical meaning. We therefore adopt the Caputo derivative, defined as [15–17]:

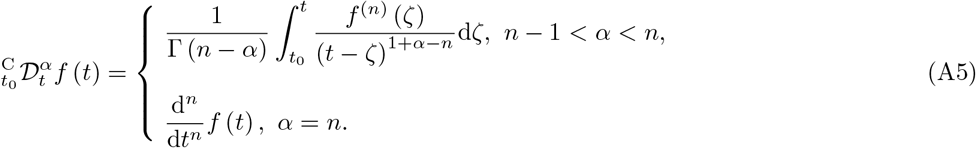

Here, 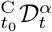 denotes the Caputo fractional derivative. The superscript C denotes the Caputo form, *t*_0_ is the initial time (set to *t*_0_ = 0), and *t* is the current time. Specifically, when *α* ∈ (0, 1), Eq. (A5) simplifies to:

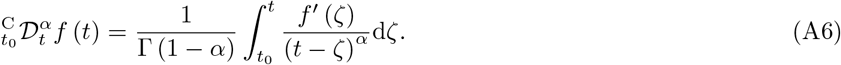

For notational simplicity, we abbreviate 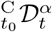 as 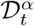 throughout this work. It can be seen from Eq. (A5) and Eq. (A6) that past influence follows a power-law decay (Fig. S1).

Let 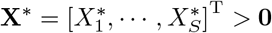 be a feasible equilibrium such that 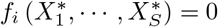. Because infeasible equilibria imply species extinction, stability analyses typically focus on feasible equilibria. Near such equilibria, the dynamics of a memory system can be approximated by:

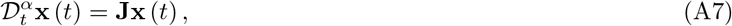

where **x** (*t*) = **X** (*t*) − **X**^*^ is the perturbation and **J** is the community matrix, with *J*_*ij*_ capturing species *j*’s influence on species *i* near equilibrium. The equilibrium is stable if lim_*t*→+∞_ **x** (*t*) = **0**.

For memoryless systems, Eq. (A7) reduces to:

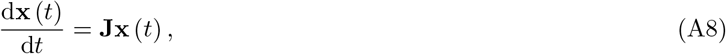

whose solution, given initial perturbation **x**_0_ at *t* = 0, is:

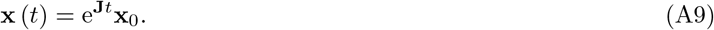

To ensure **x**(*t*) → **0** as *t* → ∞, all eigenvalues *λ* of **J** must satisfy:

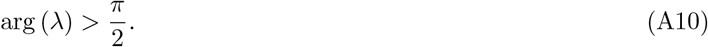

Geometrically, this implies that eigenvalues must lie in the left half of the complex plane.

For systems with memory, solutions to Eq. (A7) involve the Mittag-Leffler function *E*_*α*_(*z*), defined as [44]:

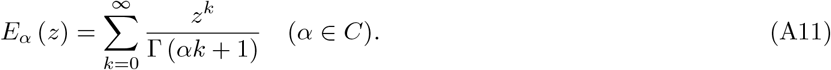

When *α* = 1, it reduces to the exponential function:

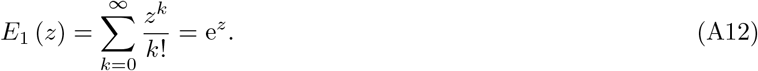

Given initial condition **x**_0_, the solution becomes:

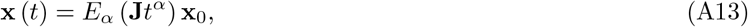

or in expanded eigenvalue form:

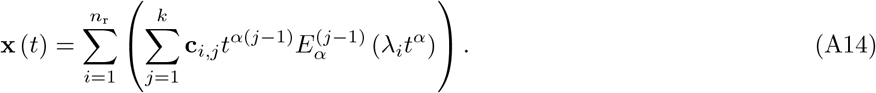

Here, *n*_r_ is the number of eigenvalues, *k* is the algebraic multiplicity, and 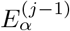 is the (*j* − 1)-th derivative.

Asymptotic properties of the Mittag-Leffler function include [44]:

1. If |arg(*s*)| ≤ *απ/*2, then

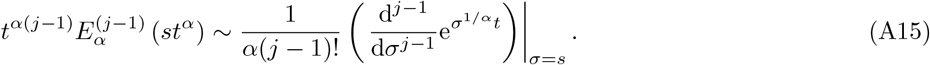
2. If |arg(*s*)| ≤ *απ/*2, then

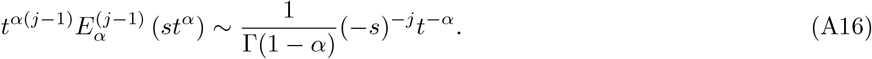

These properties imply that stability is guaranteed when all eigenvalues satisfy:

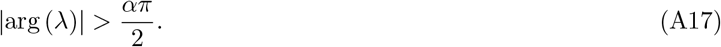

Geometrically, the instability region shrinks to a cone in the right half-plane, indicating that memory effects expand the stability region. This expansion is especially significant when instability in memoryless systems arises from conjugate eigenvalues near the imaginary axis.

### 2. Quantifying the size of the stability region for systems with memory

Based on the stability criterion for systems with memory, the angle of the instability sector in the complex plane is given by

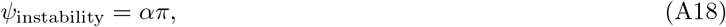

and thus the angle corresponding to the stability region becomes

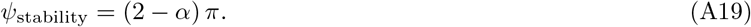

The relative size of the stability region can be quantified by the ratio between the angle of the stability region and the full angle of the complex plane (2*π*), leading to:

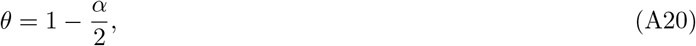

where *θ* denotes the normalized size of the stability region.

By substituting the relationship between the differentiation order *α* and memory strength ℳ into Eq. (A20), we obtain:

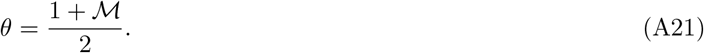

This formulation reveals that when there is no memory (i.e., ℳ = 0), the stability region corresponds to the left half of the complex plane, yielding *θ* = 1*/*2. As memory strength increases, *θ* increases accordingly, indicating an expansion of the stability region. This pattern further implies that, in systems where instability arises from a pair of complex conjugate eigenvalues crossing the instability boundary, increasing memory strength can exert a substantial stabilizing effect.

## Appendix B: Stability analysis of structured food web with memory

Following the tradition initiated by May and adopted by many subsequent researchers [3–6, 8, 9, 35–37, 40–42], we model ecological systems by directly constructing their community matrices using the random matrix modeling approach. To incorporate the trophic structure of real-world food webs, we employ the widely used cascade model [4, 5, 21, 22]. In this model, species are arranged in a strict hierarchy (i.e., trophic levels), with higher-ranked species preying on lower-ranked ones with a fixed probability.

As shown in our previous analysis, the eigenvalue distribution of the community matrix plays a central role in determining the stability of complex ecological systems. We therefore begin by presenting the analytical eigenvalue distribution for the systems studied in this work. Based on this distribution, we then perform the corresponding stability analysis.

### 1. The eigenvalue distribution of the community matrix

Since the elements of the community matrix are randomly sampled, the statistical properties of the community matrix play an important role in determining its eigenvalue distribution, as suggested by random matrix theory. Therefore, we first give these statistical properties. For upper triangular part, the mean and variance can be obtained as

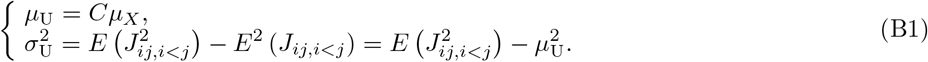

For lower triangular part, the mean and variance can be obtained as

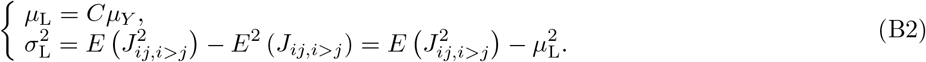

Since 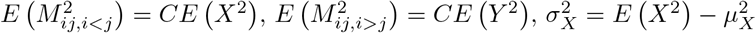, and 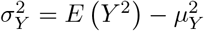, the variances of the upper triangular and lower triangular parts can be drawn as

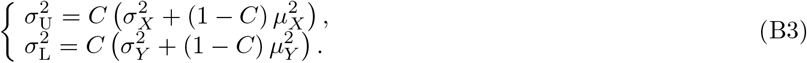

The covariance of pairwise elements of the community matrix (*J*_*ij*_, *J*_*ji*_) can be calculated as

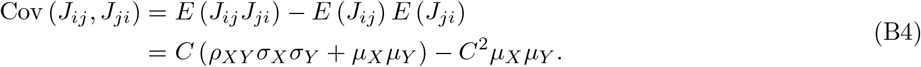

In short, statistical properties of the community matrix can be summarized as follows

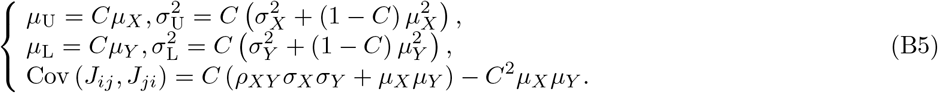

Since all diagonal elements of the community matrix are the same, the influence of having these elements is to shift the whole eigenvalue distribution to the left by *s* units. Therefore, we here consider the eigenvalue distribution of matrix 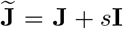. Matrix 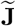 can be decomposed as 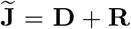, where **D** is a deterministic matrix whose upper triangular elements are all equal to *µ*_U_ while lower triangular elements are all equal to *µ*_L_. **R** is obtained accordingly as 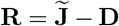. The eigenvalue distribution of 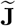 can be obtained by combining the eigenvalue distributions of **D** and **R** [5].

#### a. The eigenvalue distribution of D

For the deterministic matrix **D**, we first consider the eigenvalue distribution of the following matrix

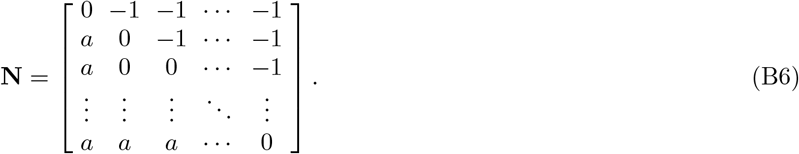

Eigenvalues *λ*_**N**_ of matrix **N** are determined by following characteristic equation

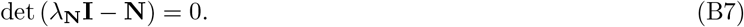

It can be shown that

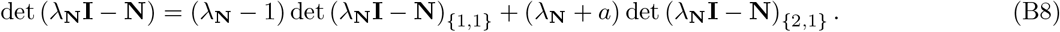

Here det (*λ*_**N**_**I** − **N**)_{*i,j*}_ represents the determinant of the matrix obtained by removing the *i*-th row and the *j*-th column from *λ*_**N**_**I** − **N**. Denote *D*_*S*_ = det (*λ*_**N**_**I** − **N**), we will have *D*_*S*−1_ = det (*λ*_**N**_**I** − **N**)_{1,1}_. Since det (*λ*_**N**_**I** − **N**)_{2,1}_ = (*λ*_**N**_ + *a*)^*S*−2^, det (*λ*_**N**_**I** − **N**) can be represented as

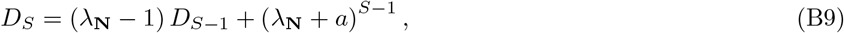

which leads to

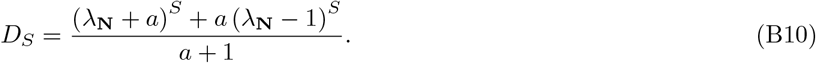

The characteristic equation (Eq. (B7)) now becomes

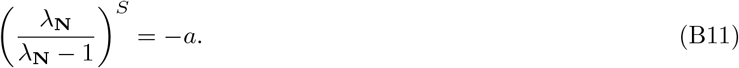

The eigenvalues of matrix **N** can then be obtained as

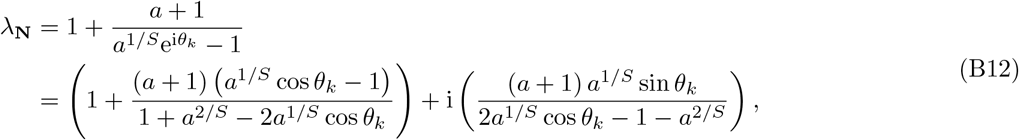

where *θ*_*k*_ = *π* (2*k* − 1)/ *S, k* = 1, 2, · · ·, *S*.

It can also be proved that the eigenvalues of **N** are actually located on a circle (note that not the full circle) centered at (*c*_**N**_, 0). *c*_**N**_ and the radius of this circle *r*_**N**_ are

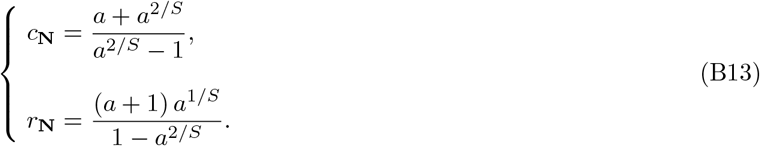

The proof can be as follows. Let

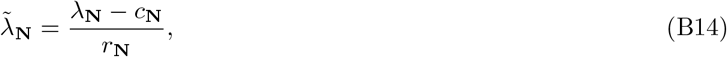

this leads to

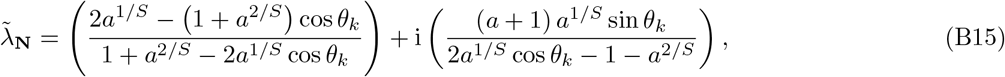

where *θ*_*k*_ = *π* (2*k* − 1)/ *S, k* = 1, 2, · · ·, *S*. It can be calculated that

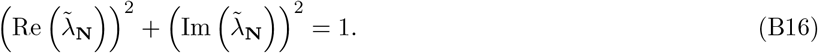

This proves that all 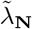 are distributed on a unit circle. Therefore, the eigenvalues of **N** are located on a circle with center (*c*_**N**_, 0) and radius *r*_**N**_.

We can then focus on the eigenvalue distribution of the deterministic matrix **D. D** can be written as

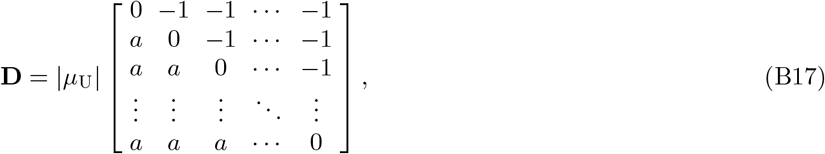

where *a* = −*µ*_L_*/µ*_U_. Therefore, the eigenvalues of matrix **D** can be derived as

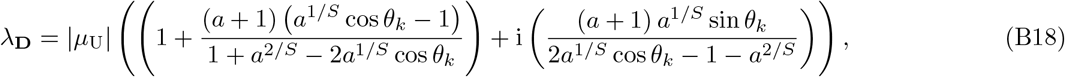

where

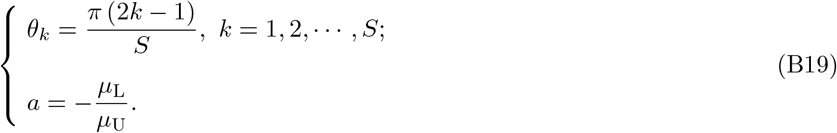

Also, the eigenvalues of **D** are located on a circle (again, not the full circle) with center (*c*_**D**_, 0) and radius *r*_**D**_, where

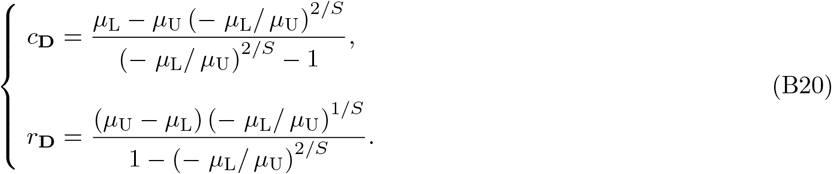

#### b. The eigenvalue distribution of R

Since matrix **R** is obtained through the difference 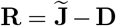, this matrix **R** can be regarded as a matrix constructed by sampling the pairs of coefficients from a bivariate distribution with means [0, 0]^T^ and covariance matrix

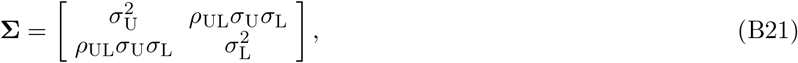

where the correlation *ρ*_UL_ can be calculated as

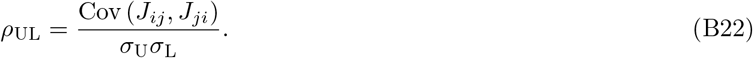

To obtain the eigenvalue distribution of matrix **R**, following assumptions are introduced [5]:

1. When *S* is sufficiently large, the eigenvalues of **R** are approximately and uniformly distributed in an ellipse centered at (0, 0), and with semi horizontal axis *r*_**R**,h_ and semi vertical axis *r*_**R**,v_;
2. The sum of these two semi axes, i.e., *r*_**R**,h_ + *r*_**R**,v_, only depends on the variances and size of the matrix, not on *ρ*_UL_;
3. The square of each semi-axis is a second-order polynomial in *ρ*_UL_, i.e., 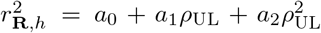 and 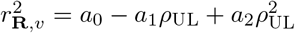.

Actually, following theoretical analysis and numerical simulations will show that these three assumptions are reasonable.

The problem now becomes the identification of the parameter *a*_0_, *a*_1_ and *a*_2_. To do so, we first consider the variances of the eigenvalues of **R**. Since tr (**R**) = ∑ *λ*_**R**,*i*_ = 0, the mean of the eigenvalues of **R** can be derived: *E* (*λ*_**R**_) = 0. Thus, the variance of the eigenvalues can be expressed as

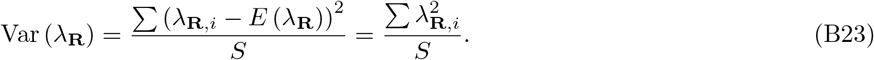

This variance can also be calculated through second-order moment

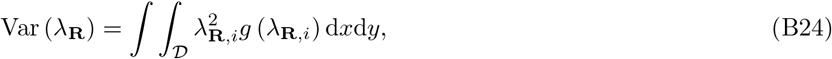

where 𝒟 is the eigenvalue distribution region in the complex plane, *g* (·) denotes the density function of the eigenvalue distribution. Since it is already assumed that the eigenvalues are uniformly distributed in an elliptic region, *g* (·) is a constant and equals 1*/πr*_**R**,h_*r*_**R**,v_. Rewrite *λ*_**R**,*i*_ as *x* + i*y*, Eq. (B24) can be further written as

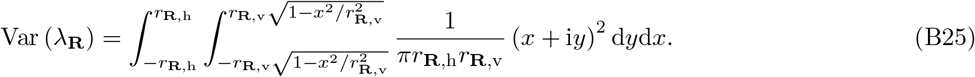

Note that the eigenvalues of **R** are either real numbers or complex conjugate pairs. Therefore, for each *λ*_1_ = *x* + i*y*, there exists *λ*_2_ = *x* − i*y*. We can then have 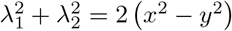. As a result, Eq. (B25) can be calculated as

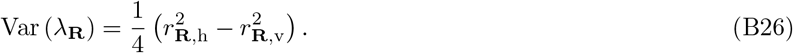

In the meantime, Var (*λ*_**R**_) can also be obtained in the following way. The fact that *λ*_**R**,*i*_ is an eigenvalue of **R** suggests that 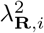 is an eigenvalue of **R**. Thus, we have tr 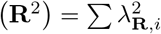. Since the *i*-th diagonal element of **R**^2^ 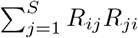 it can thus be estimated as (*S* − 1) *E* (*R*_*ij*_*R*_*ji*_). Moreover, *ρ*_UL_ is defined as

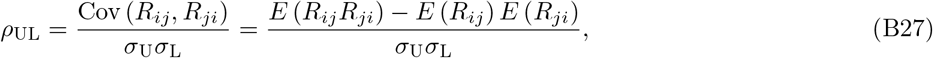

this leads to *E* (*R*_*ij*_*R*_*ji*_) = *ρ*_UL_*σ*_U_*σ*_L_. Thus, Var (*λ*_**R**_) can be obtained

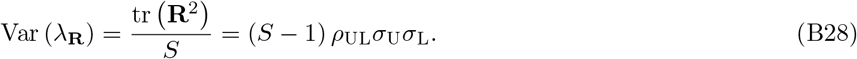

The combination of Eq. (B26) and Eq. (B28) leads to

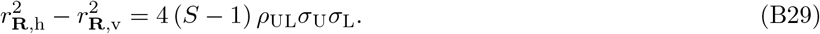

Since we already have the assumption of the exact form of *r*_**R**,h_ and *r*_**R**,v_, the parameter *a*_1_ can then be solved as

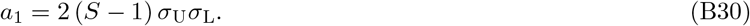

According to the assumptions introduced above, the sum of *r*_**R**,h_ and *r*_**R**,v_ does not depend on *ρ*_UL_. Therefore, *r*_**R**,h_ + *r*_**R**,v_ will be the same whenever *ρ*_UL_ = 0 or *ρ*_UL_ = 1. If *ρ*_UL_ = 0, we have

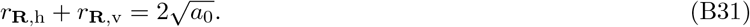

If *ρ*_UL_ = 1, we have

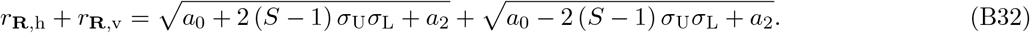

Since Eq. (B31) and Eq. (B32) should yield the same value, the value of *a*_2_ can be solved in terms of *a*_0_

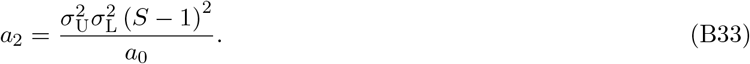

This leads to

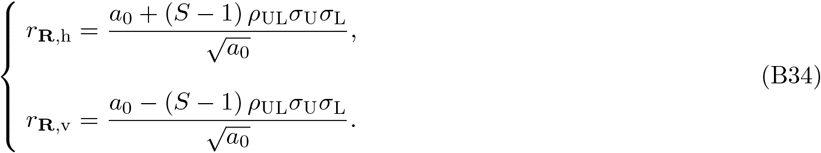

Results from Eq. (B34) suggest that the problem has now been simplified to solve for *a*_0_. Since *a*_0_ is independent of *ρ*_UL_, one can consider the case of *ρ*_UL_ = 1 to simplify analysis. Recent advances in random matrix theory suggests that *a*_0_ can be approximated as *a*_0_ ≈ *λ*_**G**,1_, where *λ*_**G**,1_ is the largest eigenvalue of the deterministic matrix

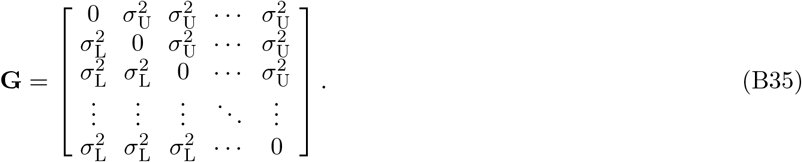

For simplicity, we can divide each element of **G** by 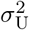 and define 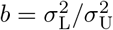, leading to the matrix

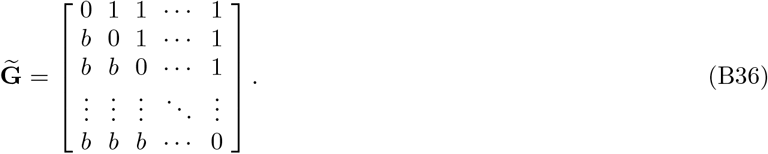

Previous analysis on the eigenvalue distribution of deterministic matrix **D** suggests that the characteristic equation of matrix 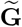 can be written as

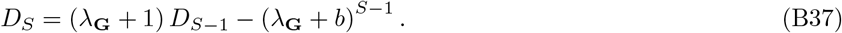

By solving this equation, we can further obtain

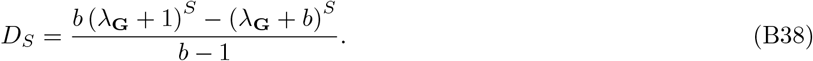

Let *D*_*S*_ = 0, we can obtain the largest eigenvalue of 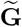

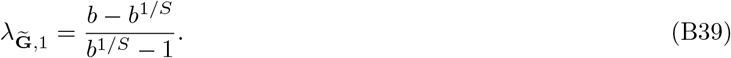

By taking the limit of 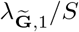 for *S* → +∞, we have

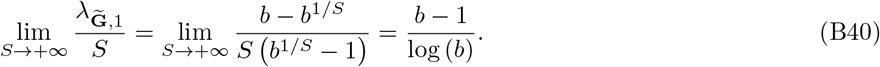

Therefore, we can get the value of parameter *a*_0_

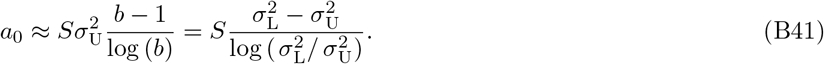

In summary, the lengths of the semi-horizontal axis and semi-vertical axis are

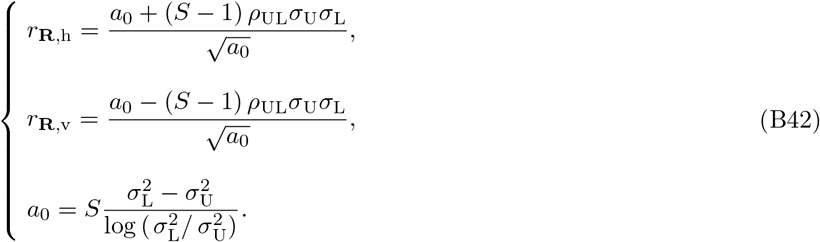

#### c. The eigenvalue distribution of J

The eigenvalues of the matrix 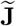 can be divided into two parts: eigenvalues located on a circle, and eigenvalues located within an ellipse. Previous studies suggest that, the first part can be approximated by the eigenvalue distribution of **D**, while the second part can be approximated by the eigenvalue distribution of matrix **R**, but with a different center ((*c*_**D**_, 0) instead of (0, 0)). Note that several eigenvalues from the first part also fall within the ellipse. By introducing the effect of diagonal elements, the eigenvalue distribution of the community matrix **J** can be obtained as follows (Fig. S2)

1. Eigenvalues located on a circle. These eigenvalues can be obtained as

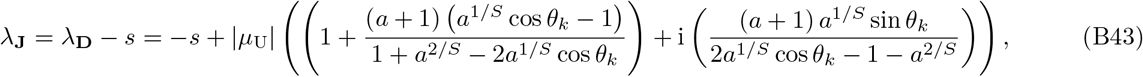

where

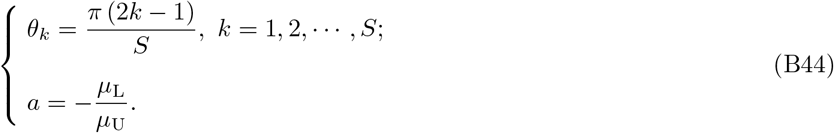
2. Eigenvalues located within an ellipse. The distribution region for these eigenvalues are

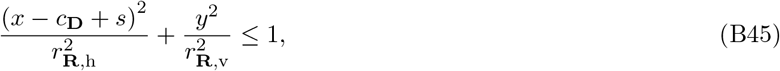

where

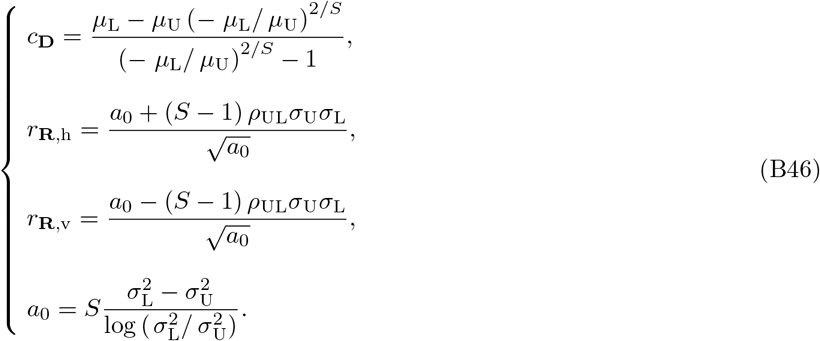

### 2. Stability analysis of structured food webs with memory

With the theoretical eigenvalue distribution of the community matrix identified, we can now utilize the general stability criterion (Eq. (A17)) to analytically determine the stability of structured food webs with memory. As suggested by our previous analysis, the introduction of memory effect expands the stability region: the instability region now becomes a sector area in the right half of the complex plane, instead of the whole right half of the complex plane. A direct observation from this stability region change is that, systems whose instability is triggered by a pair of conjugate eigenvalues crossing the imaginary axis that can be significantly stabilized by memory effect. Then, we are wondering what kind of structured food webs can be significantly stabilized by memory effect.

For memoryless systems, stability is determined by the rightmost eigenvalue of the community matrix **J**. Focusing on the structured food webs discussed in our work, as is suggested above, the eigenvalues of the community matrix can be divided into two parts: those on a circle (denoted as *λ*_**J**,c_), and those contained within an ellipse (denoted as *λ*_**J**,e_, note that several eigenvalues from the first part can also be contained in this ellipse).

For the fist part, the rightmost among these eigenvalues strongly depends on the magnitudes of *µ*_*X*_ and *µ*_*Y*_ (remember that *µ*_*X*_ *<* 0 and *µ*_*Y*_ *>* 0). If | *µ*_*X*_ | ≥ |*µ*_*Y*_ | (i.e., negative effects outweigh positive effects), the rightmost eigenvalue is a real number, and is

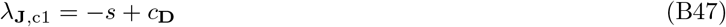

(Fig. S2). Note that this eigenvalue − *s* + *c*_**D**_ falls in the ellipse. If | *µ*_*X*_ | *<* | *µ*_*Y*_ | (i.e., negative effects are outweighed by positive effects), the rightmost eigenvalues are a pair of conjugate eigenvalues, and these two eigenvalues are

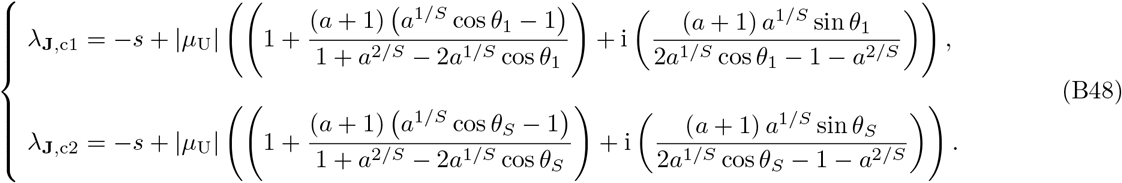

(Fig. S2). For the second part, the rightmost eigenvalue is the right endpoint of the ellipse, which is

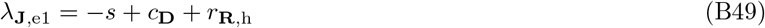

(Fig. S2).

Combining results from numerical simulations (Fig. S2), we find that when | *µ*_*Y*_ | is significantly greater than | *µ*_*X*_ | (i.e., positive effects significantly outweigh negative effects), rightmost eigenvalues are *λ*_**J**,c1_ and *λ*_**J**,c2_, otherwise rightmost eigenvalue is *λ*_**J**,e1_. Therefore, structured food webs can be stabilized by memory effect when positive effects significantly outweigh negative effects, while structured food webs where negative effects outweigh positive effects can not be stabilized by memory effect. Suppose self-regulation strength is used as the control parameter. For communities where positive effects outweigh negative ones, those with memory require a lower level of self-regulation to ensure stability, while those without memory require a higher level of self-regulation. For other communities, those with memory require similar level of self-regulation as those without memory. Similar patterns can also be observed when other community parameters are chosen as the control parameter.

Apart from identifying which type of food webs can be stabilized, we further find that the stabilizing influence of memory effect has an upper limit for these structured food webs. That is, as memory strength increases, the stabilizing influence first gets enhanced, but plateaus once memory strength reaches a certain level.

## Appendix C: Recovery rate analysis

In the main text, short-term and long-term recovery dynamics are analyzed through measuring the distance to the equilibrium after a fixed time length *T*_perturbed_ following the perturbation. That is, suppose the system is perturbed at time *t*_0_, the distance is then measured at time point *t*_0_ + *T*_perturbed_. For short-term recovery dynamics, a relatively small *T*_perturbed_ is selected, while for long-term recovery dynamics, a relatively large *T*_perturbed_ is selected. Such analyses are primarily based on numerical simulations, we here seek to provide a theoretical foundation for these results through analyzing the initial recovery rate and long-term recovery rate for systems with memory.

In our analysis, the recovery rate (more precisely, average recovery rate) at time *t*, i.e., *κ* (*t*), is defined as follows [29, 30]

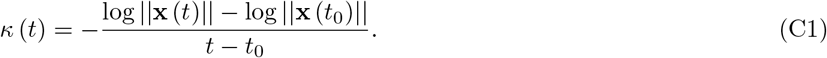

Here ||·|| is the 2-norm operator. For convenience, we suppose *t*_0_ = 0, which leads to

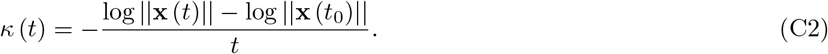

### 1. Initial recovery rate

For the convenience and simplicity of analysis, here we focus on memoryless systems where all perturbations decay initially (i.e., non-reactive systems) and their counterparts with memory, since these systems have monotonic responses to external perturbations (the distance between current system state and the equilibrium keeps decreasing).

Initial recovery rate measures the recovery rate at *t*_0_ (and in our analysis, 0), where it generally holds that

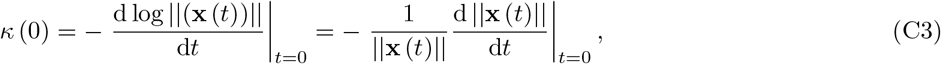

which can be further calculated as

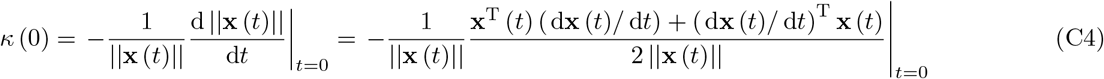

For memoryless systems, the dynamical behavior around a feasible equilibrium can be approximated by Eq. (A8), which leads to following initial recovery rate

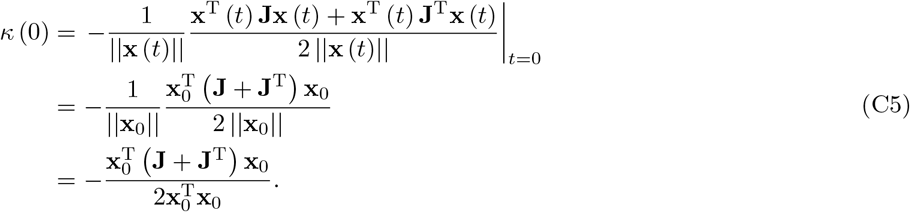

Denoting **H** = (**J** + **J**^T^) / 2, equation above becomes

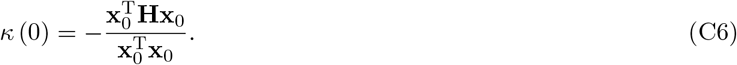

According to Rayleigh’s quotient, we have

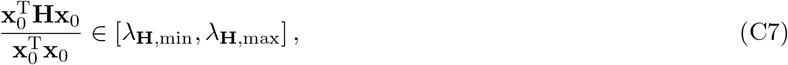

the value of *r* (0) thus falls in following range

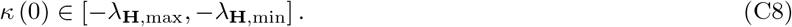

Since we are considering non-reactive systems, *λ*_**H**,max_ *<* 0. Thus, initial recovery rates of memoryless systems considered in our work are all finite positive values (Fig. S4).

For systems with memory, the dynamical behavior can be approximated by Eq. (A13), which leads to the derivative with respect to *t*

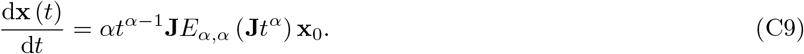

Here *E*_*α,β*_ (*z*) is the two-parameter Mittag-Leffler function, which is defined as [44]

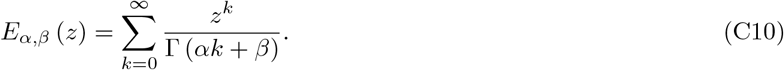

Since

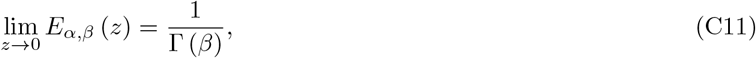

we have

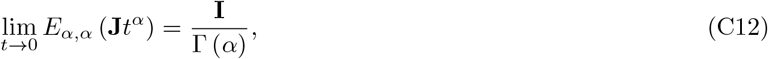

which is a finite value (**I** here represents identity matrix). Therefore, we have

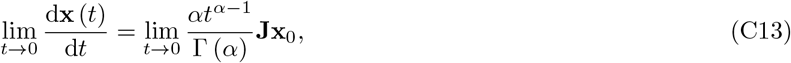

which leads to

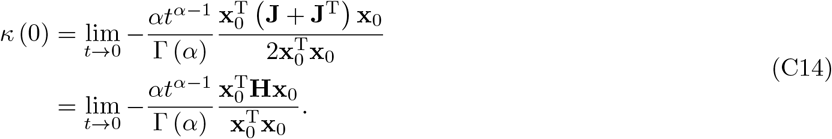

According to Eq. (C7) and the fact that we are considering non-reactive memoryless systems and their counterparts with memory, it can be proved that *r* (0) for systems with memory considered in our work shall be an infinite positive value (Fig. S4). A derivative conclusion from Eq. (C14) is that memory effects do not qualitatively influence system reactivity but quantitatively influence this property. Therefore, memory effects substantially increase initial recovery rates for these non-reactive systems. Since for non-reactive systems, the response to external perturbations is monotonic, memory effects thus greatly enhance short-term recovery.

### 2. Long-term recovery rate

In the analysis of long-term recovery rate, we only consider stable systems. For the long-term recovery rate, we assume that *t* is sufficiently large. For memoryless systems under such a condition (suppose the community matrix **J** is diagonalizable), we have

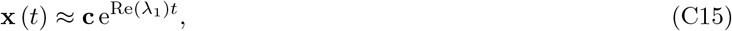

where **c** is a constant vector, and *λ*_1_ represents the dominant eigenvalue. Since we are considering stable systems, *λ*_1_ *<* 0. Therefore, we have

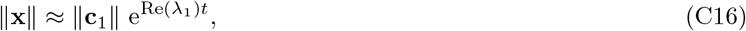

which further leads to the approximation of the long-term recovery rate:

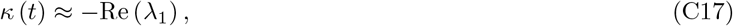

which is a constant value (Fig. S4).

Since for systems with memory, we have the asymptotic property Eq. (A16). This leads to

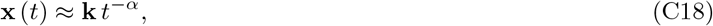

when *t* is sufficiently large. Thus, we have

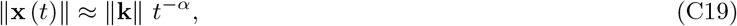

which leads to the approximation of the long-term recovery rate:

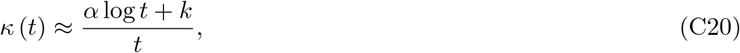

where *k* is a constant (Fig. S4).

As *t* → ∞, the long-term recovery rate of a memoryless system remains constant, while that of a system with memory approaches zero. It can thus be expected that for sufficiently large *t*, the (long-term) recovery rate of a system with memory falls below its memoryless counterpart. Therefore, we conclude that memory effects inhibit long-term recovery.

## Appendix D: Incorporating species abundance distributions

In our main analysis, we adopt the tradition initiated by May and focus directly on the community matrix **J** (i.e., the Jacobian matrix). While mathematically convenient, this tradition ignores the possible abundance distribution across different species. Here we investigate whether incorporating species abundance distributions (SADs) influences our main results. To do so, we implement the widely-used generalized Lotka-Volterra framework [1, 2]

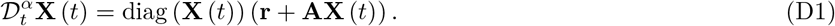

Here **r** is a *S*-dimensional vector with its component *r*_*i*_ representing the abundance of species *i*, **A** is the interaction matrix with its element *A*_*ij*_ representing the per capita influence that species *j* has on species *i*. With this framework, the role played by SAD in structuring the community matrix **J** can be clearly seen

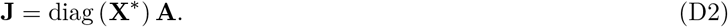

Here *S*-dimensional vector **X**^*^ is the feasible equilibrium (i.e., 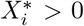 for all *i*) of the generalized Lotka-Volterra system (**r** + **AX**^*^ = **0**), and contains the information for species abundance distribution.

To investigate whether changes in SAD influence the stabilizing role of memory effects, we apply three SAD patterns in our exploitative systems: (1) Species abundances are randomly distributed across different trophic levels (i.e., uncorrelated case); (2) Species in higher trophic level tends to have higher abundance (i.e., top-abundant case); (3) Species in higher trophic level tends to have lower abundance (i.e., base-abundant case). For all three patterns, the interaction matrix **A** is constructed following the same procedure that we adopted in constructing community matrix **J**. With this constructing method, the species are already ordered according to their trophic ranks: the abundance vector **X** can be written as **X** = [*X*_1_, · · ·, *X*_*i*_, · · · *X*_*S*_]^T^, with the subscript *i* representing the trophic level. To generate all three SAD patterns, we first generate *S* random variables [*Y*_1_, · · ·, *Y*_*S*_] from the same distribution independently. For the first pattern, we keep the original order of [*Y*_1_, · · ·, *Y*_*S*_] and assign them as the equilibrium abundance for each species, i.e., 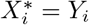. For the second pattern, we arrange these *S* variables in an ascending order 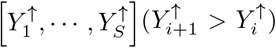 and assign them as the equilibrium abundance of each species, i.e., 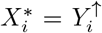. For the third pattern, we arrange these *S* variables in a descending order 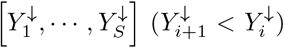 and assign them as the equilibrium abundance of each species, i.e., 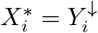.

Numerical simulations show that introducing different SAD patterns does not alter the main conclusions (Figs. S6–S8):

1. Memory effects stabilize systems with strong positive effects, but provide no stabilization for systems with either strong negative or balanced effects.
2. The stabilizing influence of memory effects is bounded.
3. Memory effects enhance short-term recovery but hinder long-term recovery.

**FIG. S1:**
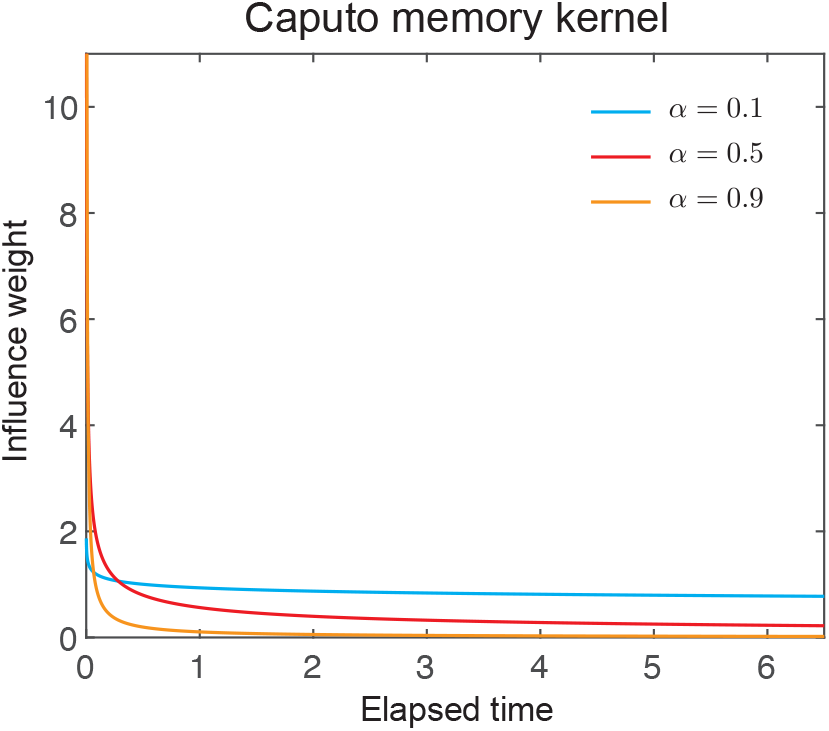
Caputo memory kernel illustrating the power-law decay of a past state’s influence. The kernel *K*_*α*_(*τ*) = Γ(1 − *α*)^−1^*τ* ^−*α*^ determines how the effect of a state that occurred *τ* units of time ago contributes to the present dynamics. Shown are kernels for three representative fractional orders (*α* = 0.1, 0.5, 0.9). Smaller *α* values correspond to slower decay, implying that past states retain influence over longer timescales, whereas larger *α* values concentrate the influence near the present. At the limit *α* → 1, the kernel collapses to a Dirac delta distribution, recovering the classical first-order derivative without memory.

**FIG. S2:**
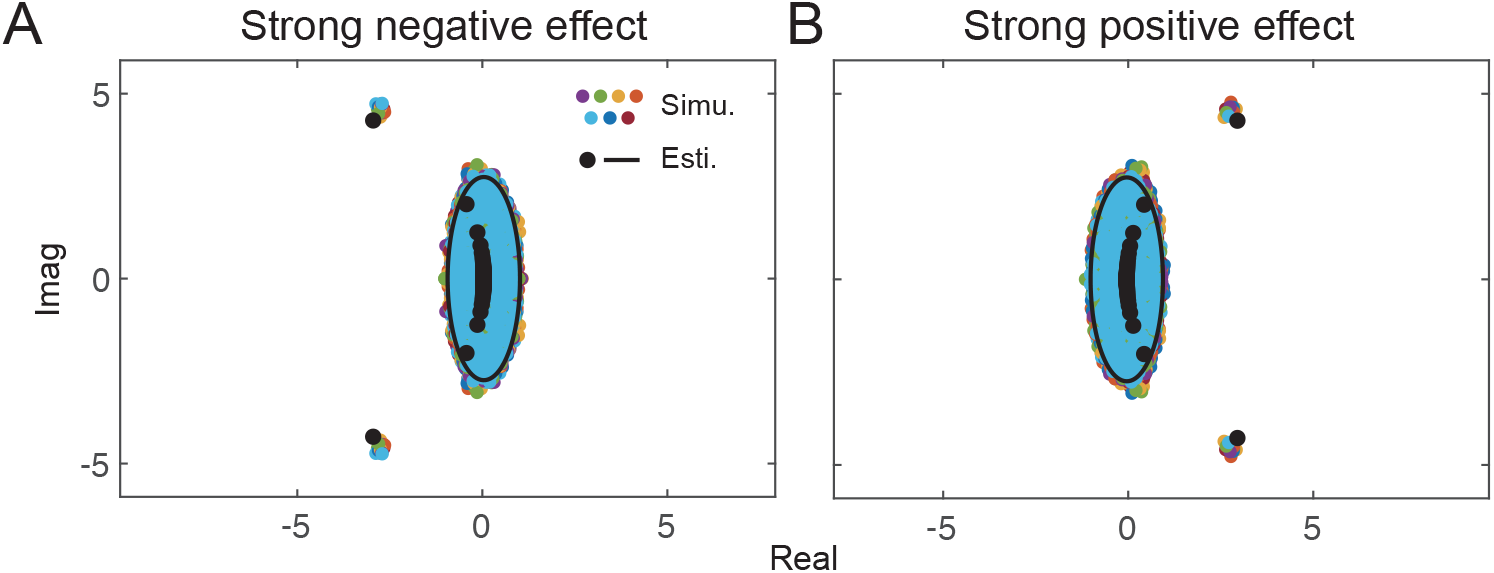
Eigenvalue distributions of the cascade community matrix. Colored dots indicate numerical simulations, while black dots and lines represent theoretical estimations. The two are in good agreement. In (A), *µ*_*X*_ = −0.9 and *µ*_*Y*_ = 0.1 (strong negative effect), and in (B), *µ*_*X*_ = −0.1 and *µ*_*Y*_ = 0.9 (strong positive effect). Other system parameters are *S* = 200, *C* = 0.1, *s* = 0, *σ* = 0.05, and *ρ* = −0.7.

**FIG. S3:**
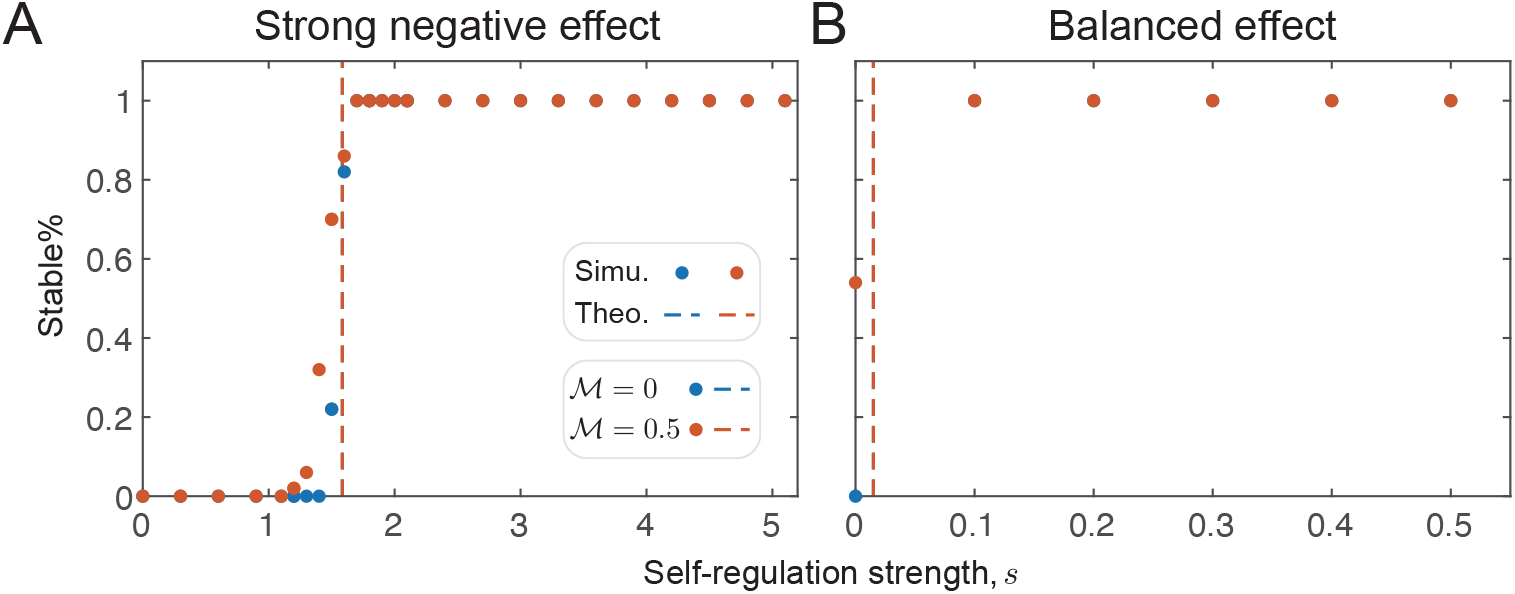
Stability of cascade exploitative systems with strong negative and balanced effects, with and without memory. Blue and red dots indicate simulation results for systems without and with memory, respectively, and dashed lines show theoretical critical thresholds. Each dot represents the proportion of stable realizations among 50 independently generated communities with the same set of system parameters. Memory does not provide stabilizing effect in either case. For strong negative effect (panel (A)), *µ*_*X*_ = − 0.9 and *µ*_*Y*_ = 0.1, and for balanced effect (panel (B)), *µ*_*X*_ = − 0.5 and *µ*_*Y*_ = 0.05. Other parameters are *S* = 500, *C* = 0.1, and *σ* = 0.05.

**FIG. S4:**
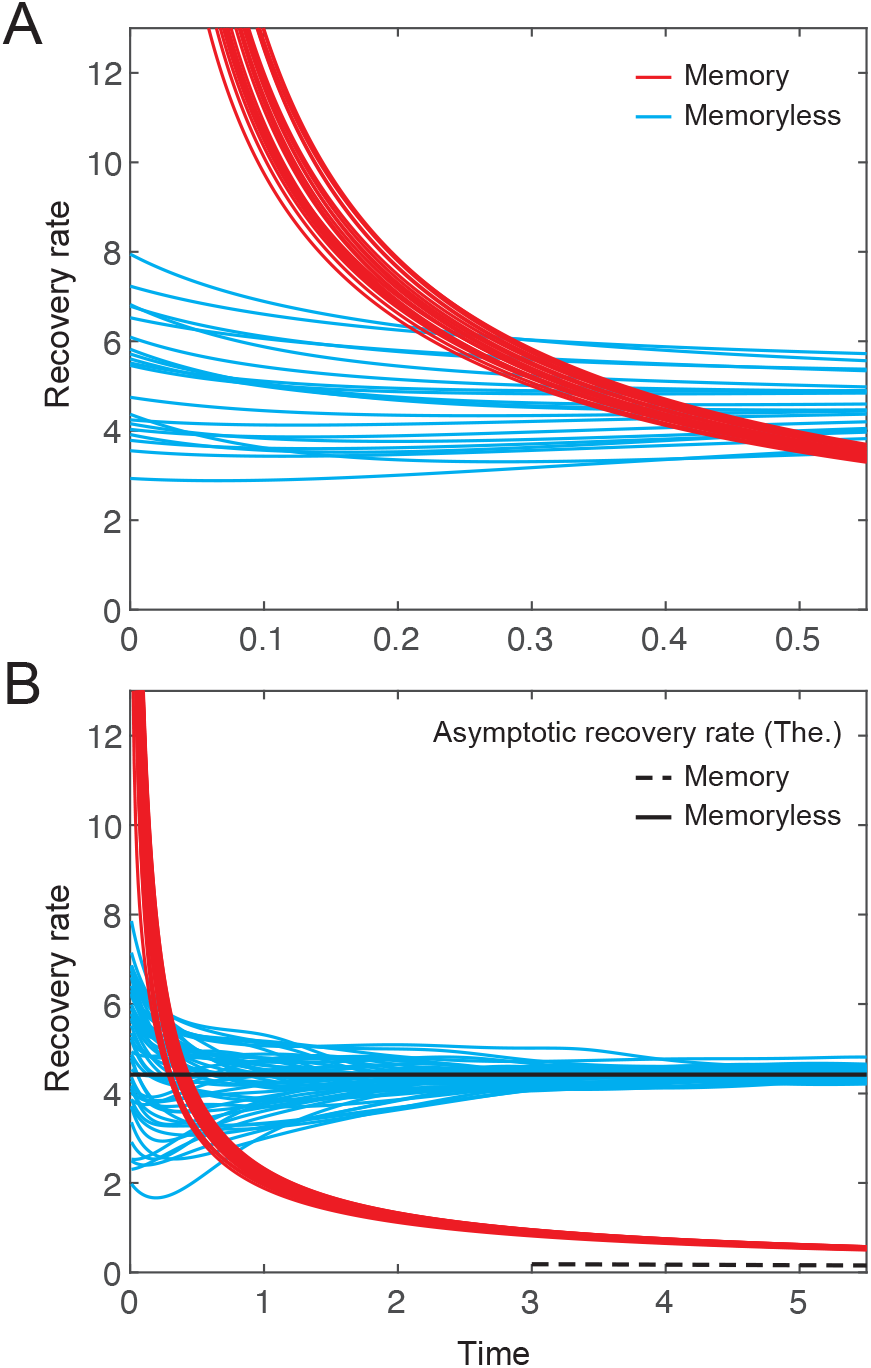
Recovery rates of systems with and without memory. Panel (A) shows short-term recovery rates following perturbations, and panel (B) shows long-term recovery rates. Blue and red curves represent systems without and with memory, respectively. Black solid and dashed lines denote theoretical asymptotic recovery rates for systems without and with memory. In each panel, recovery rates from 20 independently generated communities with the same set of system parameters are shown. Parameters are *S* = 10, *s* = 5, *C* = 0.1, *µ*_*X*_ = −0.1, *µ*_*Y*_ = 0.9, *σ* = 0.05, *ρ* = −0.7, and ℳ = 0.5.

**FIG. S5:**
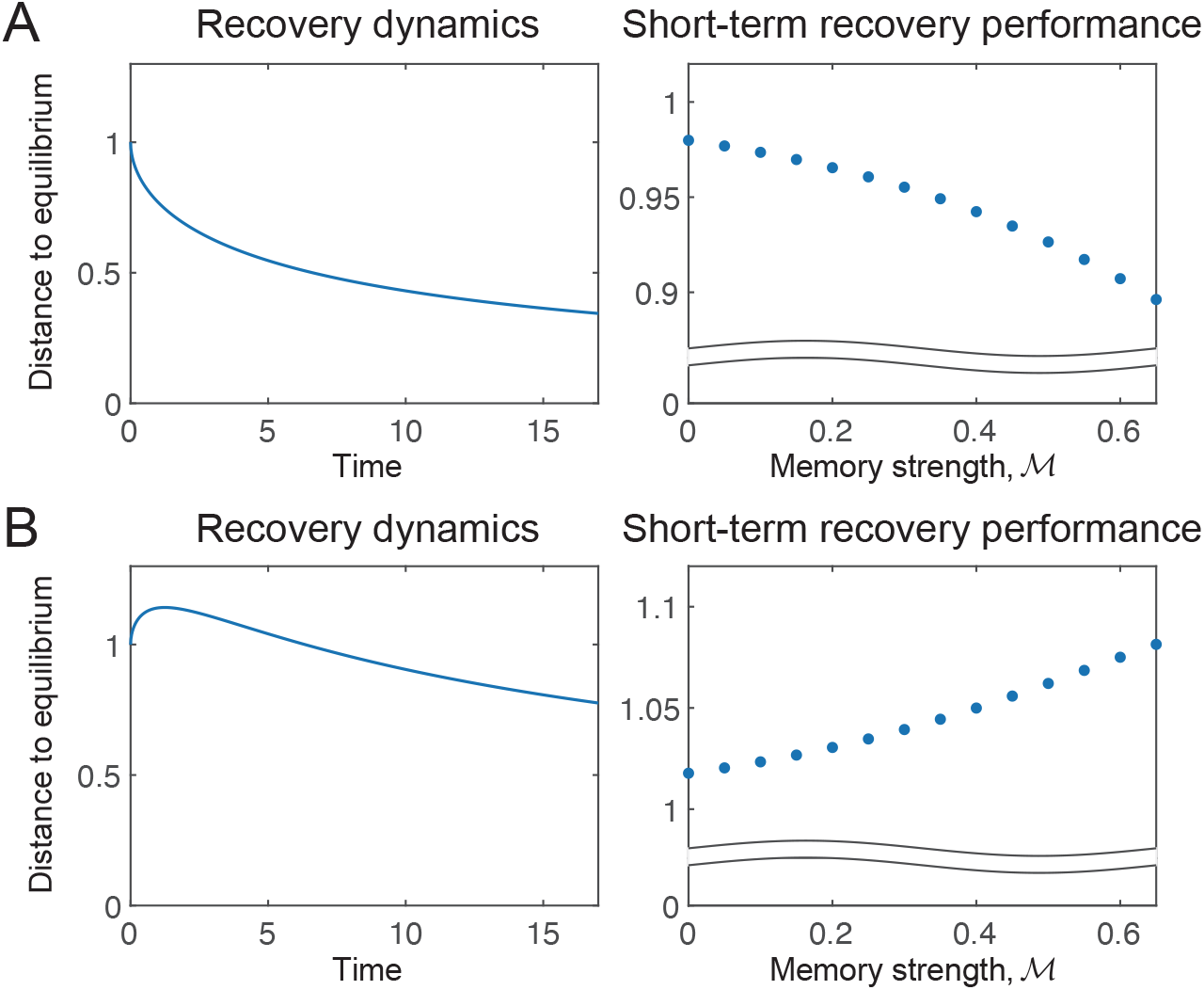
Short-term recovery of reactive systems. (A) Case where the reactive direction is not engaged during the dynamics. Left panel: recovery dynamics showing a monotonic return to equilibrium. Right panel: stronger memory consistently enhances short-term recovery. (B) Case where the reactive direction is engaged during the dynamics. Left panel: recovery dynamics showing a non-monotonic response to perturbation. Right panel: stronger memory no longer guarantees improvement in short-term recovery. All simulations are based on a two-species exploitative system modeled with generalized Lotka–Volterra dynamics. Short-term performance is evaluated at *T*_perturbed_ = 0.1. In the left panels of (A) and (B), memory strength is fixed at 0.5. Community parameters are *A*_11_ = −0.1, *A*_12_ = −0.1, *A*_21_ = 0.5, *A*_22_ = −0.1.

**FIG. S6:**
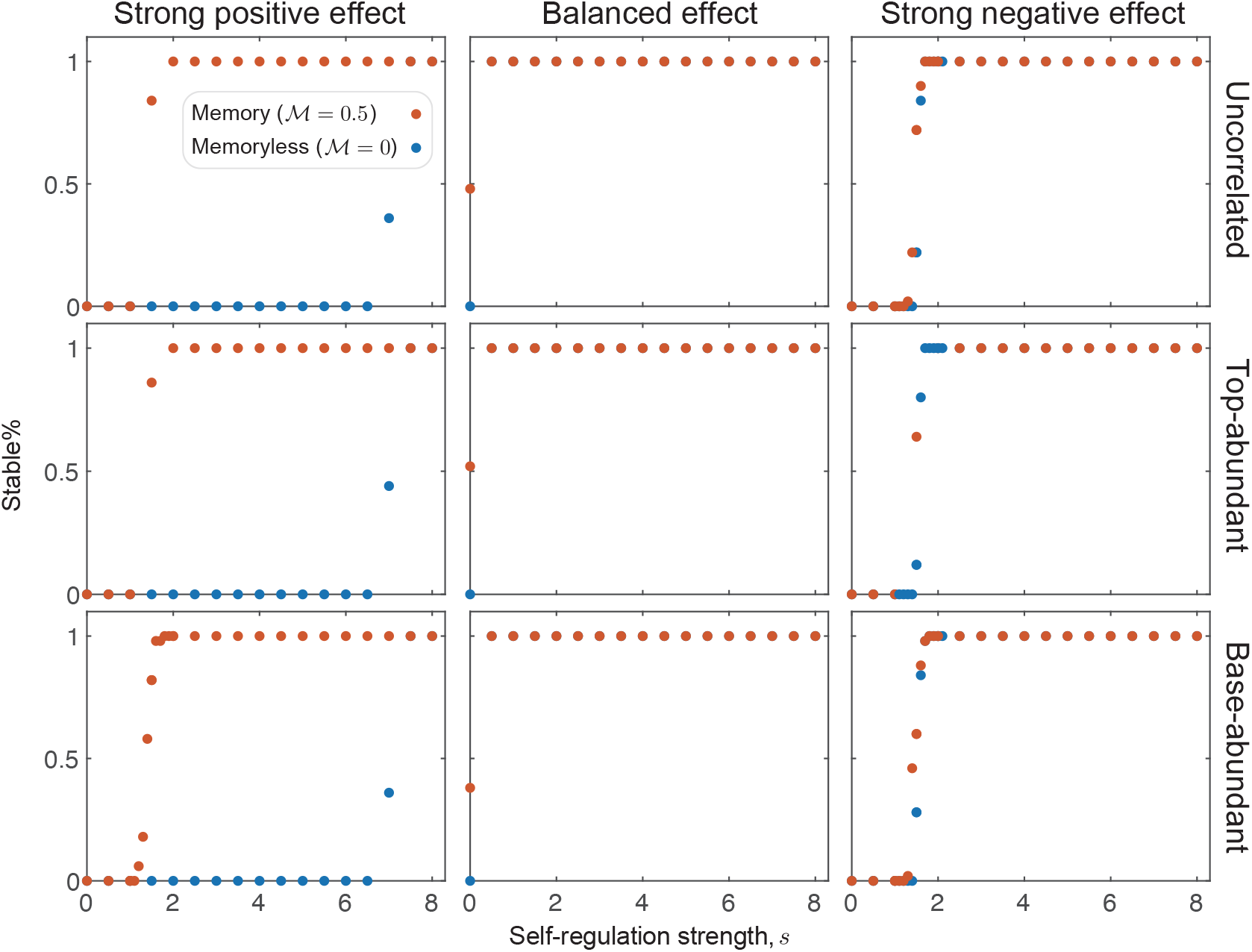
Stability of cascade exploitative systems under different species abundance distributions (SADs). Equilibrium abundances are independently drawn from a uniform distribution, with three SAD patterns considered: uncorrelated (first row), where abundances are randomly distributed across trophic levels; top-abundant (second row), where higher trophic levels tend to have higher abundances; and base-abundant (third row), where higher trophic levels tend to have lower abundances. Blue and red dots represent communities without memory and with memory, respectively. Each dot shows the fraction of stable realizations among 50 independently generated systems with identical parameters. Results indicate that memory can stabilize systems with strong positive effects, but not those with balanced or strong negative effects. System parameters: strong negative effect (first column), *µ*_*X*_ = −0.9, *µ*_*Y*_ = 0.1; balanced effect (second column), *µ*_*X*_ = −0.5, *µ*_*Y*_ = 0.5; strong positive effect (third column), *µ*_*X*_ = −0.1, *µ*_*Y*_ = 0.9. Other parameters: *S* = 500, *C* = 0.1, *σ* = 0.05, and *ρ* = −0.7.

**FIG. S7:**
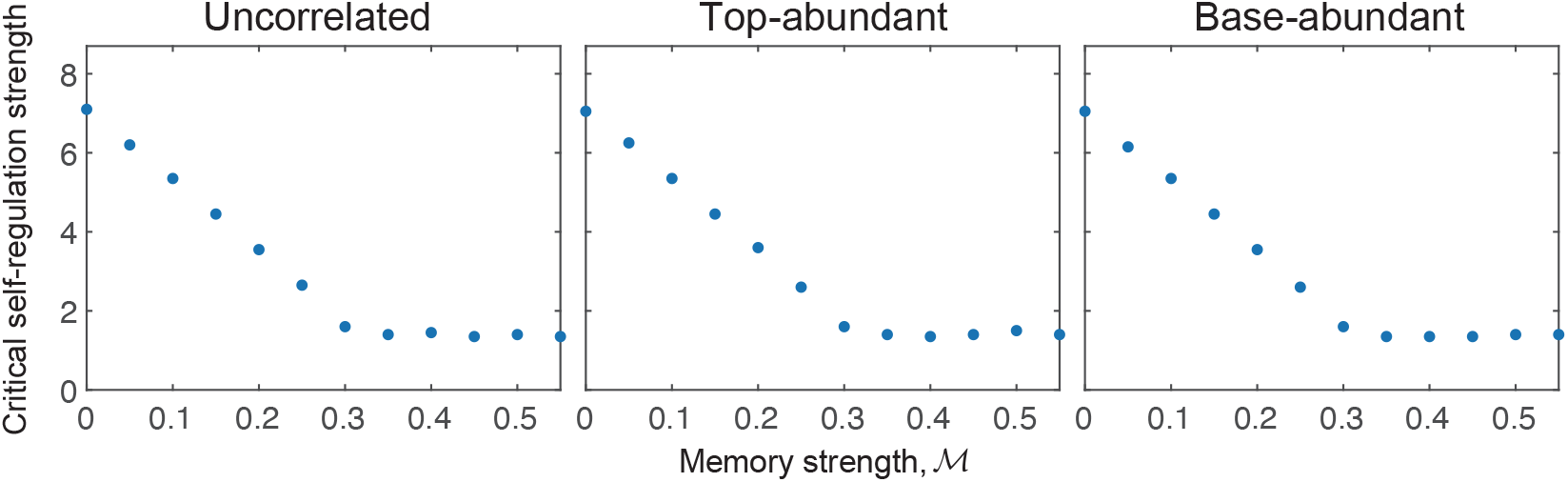
Saturating stabilizing effect of memory under different species abundance distributions. Results are shown only for the strong positive effect case, as memory has no stabilizing influence under balanced or strong negative effects. System parameters: *µ*_*X*_ = −0.1, *µ*_*Y*_ = 0.9, *S* = 500, *C* = 0.1, *σ* = 0.05, and *ρ* = −0.7.

**FIG. S8:**
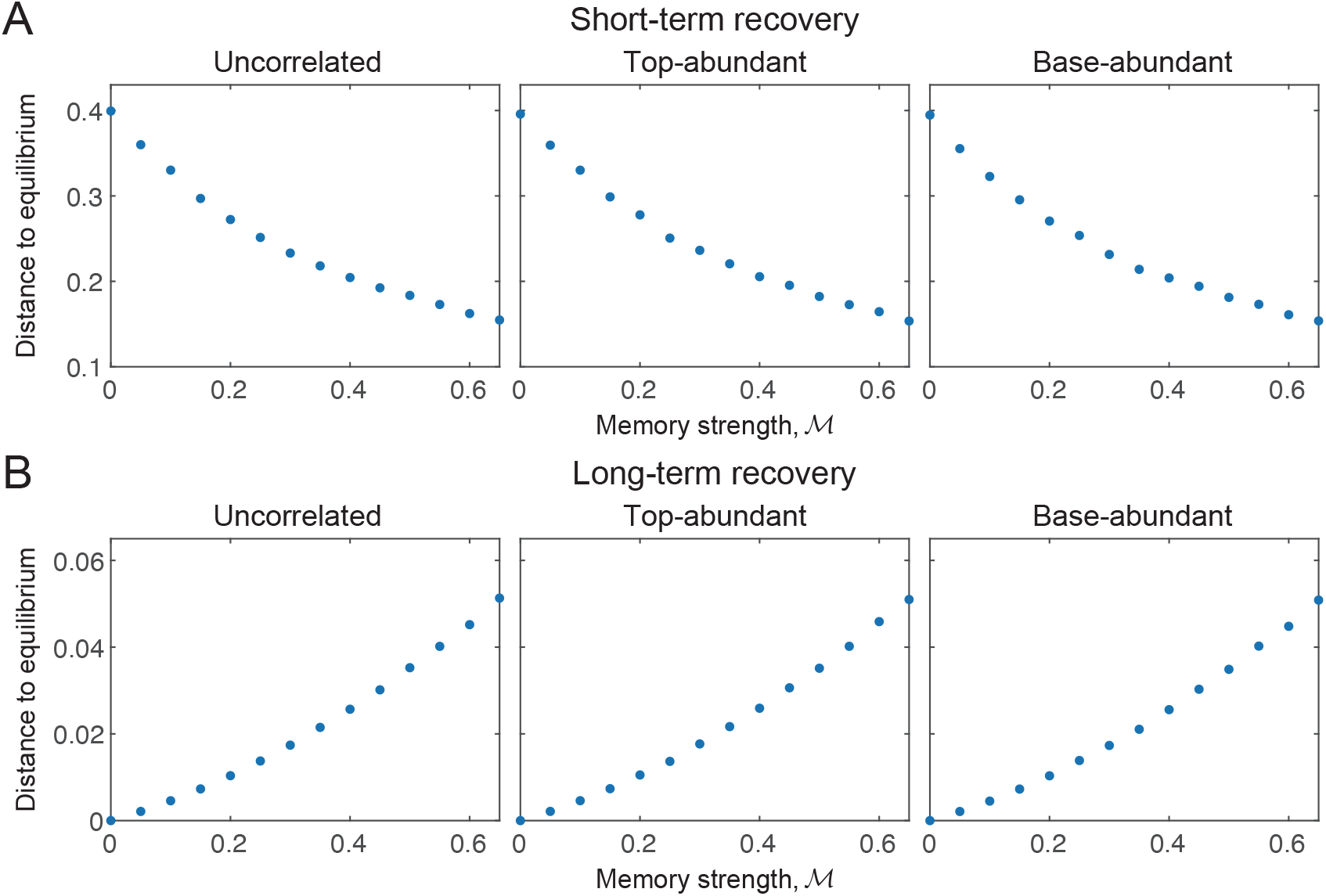
Short-term (A) and long-term (B) recovery of systems with memory under different species abundance distributions (SADs). Short-term recovery is evaluated at *T*_perturbed_ = 0.1, and long-term recovery at *T*_perturbed_ = 3. System parameters: *µ*_*X*_ = −0.1, *µ*_*Y*_ = 0.9, *S* = 500, *C* = 0.1, *σ* = 0.05, and *ρ* = −0.7.

**TABLE S1:** Summary of the main symbols used in this study.

| Symbol | Description | Note |
| --- | --- | --- |
| $X_i$ | Abundance of species $i$ | – |
| $\mathcal{D}_t^\alpha$ | Caputo-type fractional derivative operator of order $\alpha$ | – |
| $\alpha$ | Order of differentiation (memory exponent) | $\alpha \in (0, 1]$ |
| $\mathcal{M}$ | Memory strength | $\mathcal{M} = 1 - \alpha$ |
| $\mathbf{J}$ | Community matrix | – |
| $\theta$ | Stability region size | $\theta = (1 + \mathcal{M})/2$ |
| $D$ | Euclidean distance between current and equilibrium states | – |
| $s$ | Self-regulation strength | Diagonal element of $\mathbf{J}$ |
| $C$ | Connectance (fraction of realized interactions) | – |
| $\mu_X, \mu_Y$ | Mean strengths of negative and positive effects | $\mu_X < 0, \mu_Y > 0$ |
| $\sigma_X, \sigma_Y$ | Standard deviations of interaction strengths | – |
| $\rho$ | Correlation between reciprocal interactions | – |
| $\kappa$ | Recovery rate | – |

